# Elucidating the dysbiotic features of gut microbiome, interaction with the human genome, and utility as a biomarker for treatment of Parkinson’s disease

**DOI:** 10.64898/2026.05.12.724602

**Authors:** Charles F Murchison, Giacomo Antonello, Zachary D Wallen, Marissa N Dean, Adrian Verster, Kaelyn Long, Levi Waldron, Timothy R Sampson, David G Standaert, Haydeh Payami

**Affiliations:** Department of Neurology, University of Alabama at Birmingham, Birmingham, AL, 35233, USA; Aligning Science Across Parkinson’s (ASAP) Collaborative Research Network, Chevy Chase, MD, 20815, USA; Department of Biostatistics, University of Alabama at Birmingham, Birmingham, AL, 35233, USA; Graduate School of Public Health and Health Policy and Institute for Implementation Science in Population Health, City University of New York, New York, NY, USA; Pythia Informatics, Ottawa, Canada; Department of Cell Biology, Emory University School of Medicine, Atlanta GA, 30329, USA

## Abstract

Parkinson’s disease (PD) is the fastest-growing neurologic disease and a leading cause of disability worldwide. PD affects the body and mind, is progressive, and there is no prevention or cure. Gut microbiome, a recently recognized contributing factor in PD, offers new leads for understanding the underlying pathobiology and devising new treatments. Here, we present the most comprehensive study of the PD gut microbiome to date, comprising three large datasets with a sample size of 1,006 PD and 544 neurologically healthy controls, generated with uniform methodology from subject recruitment to data analysis, and characterized using deep shotgun metagenome sequencing, genome-wide genotypes, and metadata. We begin by describing the gut dysbiosis at the species, gene, pathway, and functional level. Next, we find that PD-associated genetic variants at the *SNCA* gene region are associated with increased abundance of opportunistic pathogens and depletion of fiber degraders in the PD gut. We show that the presence of opportunistic pathogens at high levels in the gut increases the penetrance of *SNCA* variants for PD risk, raising the GWAS-derived odds ratio from less than 1.5 to over 8. The genetic variants identified here control splicing of the *SNCA* transcripts into alpha-synuclein isoforms with varying affinity for pathological aggregation. These data suggest pathogens are triggers for disease in the setting of genetic susceptibility, and the link to the genome implicates the microbes in the causation of PD. Finally, shifting focus to translation, we show that not all PD patients have the same dysbiotic features, and propose a conceptual framework to identify microbiome-based biomarkers to select appropriate patients for targeted microbiome-based clinical trials and personalized treatment.

## Introduction

The prevalence of Parkinson’s disease (PD) has been increasing sharply worldwide and is expected to continue to rise ^1^. PD is a leading source of disability and death globally. Between 2000 and 2019, disability due to PD increased by 81%, and death increased by 100% ^1^. There is no currently accepted approach for prevention. Without timely and effective intervention, PD poses an escalating global public health burden ^2^.

Classically defined as a brain disorder that causes abnormal movement, PD is now recognized as a heterogeneous and complex systemic disorder ^3^. Pathology is found in the brain and in the periphery, including the gut ^4–6^, and the disease causes both motor and non-motor dysfunction ^3^. Gastrointestinal problems (constipation), sleep disorders, and anosmia can precede the onset of motor signs by decades. As the disease progresses, motor and non-motor symptoms become debilitating, and many individuals develop psychiatric problems and dementia. Treatments are symptomatic and do not slow the progression of the disease.

Except for rare highly penetrant mutations in genes like *SNCA* and *PRKN*, the causes of PD are unknown ^7^. Over 90 genetic susceptibility loci ^8^ and an array of environmental factors ^9,10^ have been associated with risk of developing PD, none of which is sufficiently penetrant to cause disease by itself. There is likely an interaction between genetic predisposition and external triggers in ways that are not yet understood. Involvement of the altered composition of the gut microbiome in PD is a more recent observation. The gut microbiome has diverse functions that are critical for human health, and imbalances in the composition of the microbiome (dysbiosis) can cause disease ^11^. Therefore, understanding the link between microbiome dysbiosis and PD could bring novel insights into disease causation and progression. Moreover, the microbiome is more amenable to manipulation and change than the human brain or genes, opening a promising new frontier for targeted treatments.

Independent lines of evidence from experimental and animal models have shown that the gut microbiome contributes to PD-relevant pathologies ^12–14^. Human studies have consistently found the gut microbiome to be dysbiotic in PD, but they have been largely inconsistent in detailing the dysbiotic features. A recent meta-analysis summarized the PD literature, including sixteen studies that used 16S amplicon sequencing (total 1367 controls, 1798 PD) and six that used shotgun metagenomics sequencing (total 770 PD, 554 controls) ^15^. They identified ∼40 taxa as being altered in PD. They make a strong point about vast study-specific differences, which hamper meta-analysis and limit generalizability. They quantified the variability in the microbiome that was attributed to the study of origin as 20% for 16S and 8% for shotgun studies, as compared to 1% that is attributed to PD vs. controls. The major biological effectors of microbiome (age, sex, body mass index, diet, and Bristol stool chart (constipation)) each account for less than 5% of variability in gut microbiome ^16^, which further underscores the striking 8%-20% variation stemming from differences in studies. Inter-study variation, exacerbated by the lack of standardized methodology, has been a major obstacle in the field, causing low concordance across studies and draining power in meta-analyses. Techniques are available to harmonize raw sequences from different sources for bioinformatic and statistical meta-analysis, but little can be done for upstream differences in how samples were collected, stored, DNA extracted, and sequencing was performed.

We present a study of three datasets from across the United States (US), which were collected and analyzed using uniform methodologies to the degree possible. The sample size is larger than all shotgun studies to date combined, consisting of 1,006 PD and 544 neurologically healthy controls (referred to as NHC or controls). With this large sample size and power not lost to inter-study variation, we conducted a three-pronged study. First, we delineated the dysbiotic features of the PD gut microbiome, conducting direct replication and meta-analysis. Second, we tested for interaction between alpha-synuclein gene (*SNCA*) variants and dysbiotic features of PD microbiome. *SNCA* is the most important gene for PD; it encodes a protein that aggregates, forming Lewy bodies and neurites, which are the pathological hallmarks of PD. Highly penetrant mutations in *SNCA* cause autosomal dominant PD. Low-penetrance non-coding variants in and around *SNCA* are associated with increased risk of idiopathic PD and are the highest peak in genome-wide association studies (GWAS) of PD ^8^. The genetic effect size (penetrance), however, is small, leaving room for as-yet unidentified triggers and modifiers.

Here, we show interaction between *SNCA* genotype and the relative abundance of opportunistic pathogens and fiber-degrading microbes and provide evidence that opportunistic pathogens elevate penetrance of *SNCA* risk genotype several-fold. Pivoting to translation, we investigate heterogeneity and subtypes of PD at the microbiome level and explore the feasibility of using dysbiotic features as biomarkers and diagnostic companions. PD is highly heterogeneous. One drug will not work for all patients, and in fact, clinical trials for disease-modifying treatments have not succeeded, partly because the heterogeneity in cohorts dilutes any potential efficacy signal. Biomarkers that would enable subtyping patients to select the appropriate cohort for a given drug are a high-priority and unmet need in PD. With the increasing interest in targeting the microbiome for drug development and treatment, we present a framework for biomarker development as diagnostic companions, exploring ways to empower clinical trials and venture into potential uses in personalized treatments.

We have generated new large datasets, linking shotgun metagenomes with genome-wide genotypes and metadata, and share the data publicly without restriction. We provide both raw and processed data at the individual level (see **Data and code availability** Section). The raw data can be processed and analyzed in new ways. The processed and analyzed data are provided in their entirety in user-friendly Excel files as **Extended Data,** where readers can search and identify features of interest if not explicitly reported in the manuscript.

## Results and discussion

The following text, main figures, and tables are summaries and syntheses of results as presented in their entirety in **Extended Data**. **Extended Data** files are organized by topic into eight Excel files, each with multiple tabs starting with a README that outlines and lays out the rationale for the data contained in each tab. **Extended Data** can be reproduced using the raw data and analytic code provided, while following the **Methods** section.

The analytic sample size, after quality control and exclusions, is 1,006 persons with PD and 544 NHC (**Table 1. Participant characteristics**), larger than all published shotgun datasets of PD combined, without the confounding that results from differences in study of origin ^15^, and to our knowledge, it is the only metagenomic dataset paired with genome-wide genotypes. The study comprises three datasets from across the US, collected using uniform methodologies (see the **Standardization** in **Methods** section). The workflow, from subject enrollment to the generation and processing of datasets, is shown in the STORMS Chart following the guidelines for human microbiome research ^17^ (**Extended Data 1-STORMS**). The three datasets are called NGRC, UAB, and DOD. NGRC subjects were from the states of Washington in the northwestern US (45%), New York in the northeast (42%), and Georgia in the southeast (13%). UAB and DOD subjects are from Alabama and neighboring states in the Deep South, a culturally distinct population in the most southern regions of the US. Over 97% of subjects were white of European descent. Gut metagenomes were derived from DNA extracted from stool. Host genotypes were obtained from DNA extracted from blood or saliva. Metadata were collected using standardized questionnaires and medical records.

**Table 1.**
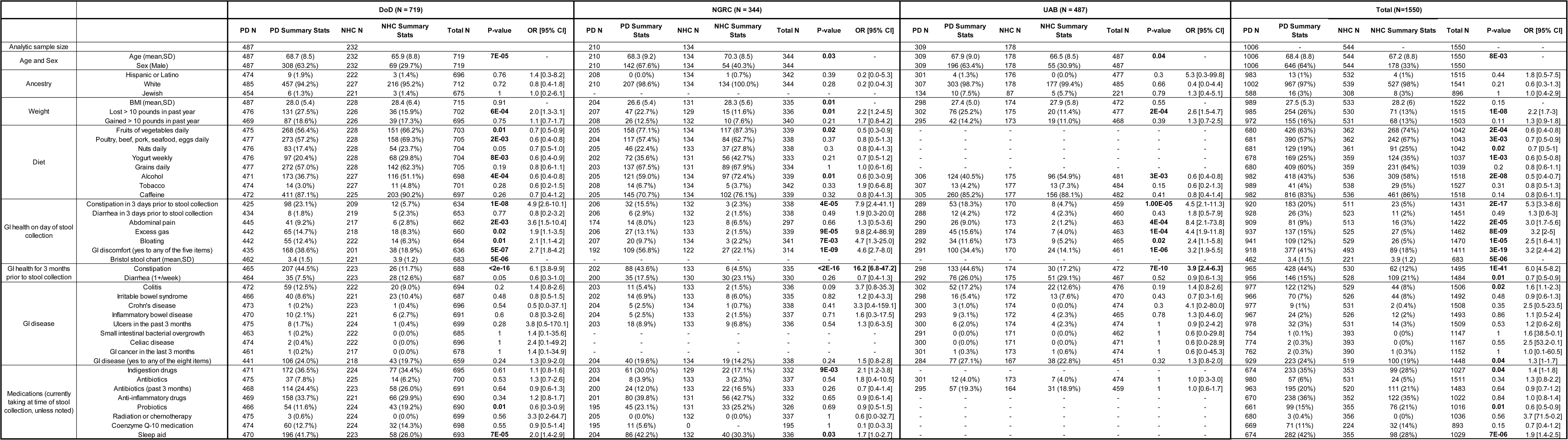
Participant characteristics. Characteristics for persons with PD and neurologically healthy controls (NHC), the difference between PD and NHC measured by odds ratio (OR) and P-value for significance (full results at individual level are in “Source data” on Zenodo).

### PD-associated features

As a validity check for metadata, we established that known PD-associated features were recapitulated in all three data sets with statistical significance (**Table 1. Participant characteristics**). As expected, more men had PD than women, in a ratio of 2 to 1. The diet of cases and controls differed; patients consuming 30% to 50% less fruit and vegetables and 40% less alcohol than NHCs. PD patients reported significant weight loss in the previous year. Patients were twice as likely to be medicated for anxiety/depression/mood and sleep aid. Most notably, GI problems, including constipation, were 4 to 16 times more common in persons with PD than NHC in every dataset (P=1E-41). Constipation is a feature of PD that can precede the onset of motor signs by decades. Later, we will address the unresolved postulate that gut dysbiosis in PD is due to constipation.

### Microbiome composition and function

Deep shotgun metagenomic sequencing of fecal DNA generated an average of 60 million sequences per subject. Sequence quality control (QC) yielded an average of 37.2 million sequences per subject which were carried into analysis. The bioBakery bioinformatics suite ^18^ was used to assign the metagenomic sequences to species, gene families, and pathways, and calculate their relative abundances.

### Microbiome-wide association studies (MWAS)

For each dataset separately (NGRC, UAB, DOD), we conducted a microbiome-wide test of association, whereby we tested differential relative abundances of species, gene families, and pathways, in turn, in PD vs. NHC (**Extended Data 2-MWAS**). Results are discussed under **Replication** and **Meta-analysis**.

### Replication across the US geography

Microbiome composition varies by geographic location ^19^, hence the concern that what appears as a dysbiotic microbiome for disease may be an artifact of regional differences. To tease out disease effects that are independent of geography, we conducted a replication study, using DOD, the larger of the two datasets from the Deep South, as discovery, and NGRC from the northern US as replication. Note that the sample size of NGRC is smaller than DOD (344 vs. 719), which can limit the power of replication. Sixty-five species that were PD-associated in DOD were detected in NGRC with reasonable prevalence (present in >5% samples) and abundance to be tested for replication.

*Escherichia coli (E. coli*) and *Klebsiella* spp were excluded (see Limitation in **Meta-analysis** section). 97% of associations trended in the same direction in NGRC as in DOD, 68% reached FDR<0.2, and 42% reached FDR<0.05 in NGRC (**Table 2. MWAS**). Among the strongest signals detected across the datasets were increased abundances in PD of *Methanobrevibacter smithii*, an archaeon that produces methane and has been linked to constipation ^20^, *Bifidobacteria* and *Lactobacillus* species which are commonly known as probiotics, opportunistic pathogens including *Porphyromonas asaccharolytica,* which are commensal to gastrointestinal and urogenital tracts but can cause infections in immunocompromised persons, *Streptococcus mutans,* an oral pathogen, and *Eisenbergiella tayi*, a pathogen that was recently shown to induce multiple sclerosis-like symptoms in germ-free mice ^21^. Bacteria that degrade plant-based fibers and produce beneficial short-chain fatty acids (SCFA), such as *Blautia wexlerae*, *Faecalibacterium prausnitzii,* and *Roseburia* species (spp), were severely reduced in PD in all three datasets. We conclude that the association of PD with microbiome is robust and reproducible, independent of geographic variation.

**Table 2.** Microbiome-wise association studies (MWAS) MWAS was conducted using Maaslin2 testing differential relative abundances of species in PD vs. NHC. Full results in Extended Data 1-MWAS. (a) Replication. MWAS in PD vs. NHC in two datasets separately, DOD representing the Deep South as discovery (N = 487 PD, 232 NHC) and NGRC representing the northern US as replication (N = 210 PD, 134 NHC). Species that reached significance (FDR<0.05) in DOD were tested for replication in NGRC. 97% trended in the same direction of fold change. Shown are species that reached FDR<0.2 in NGRC. Full results in Extended Data 1-MWAS. (b) MWAS PD vs. NHC in three datasets separately, then meta-analyzed using random-effects models (N = DOD: 487 PD, 232 NHC; NGRC: 210 PD, 134 NHC; UAB: 309 PD, 178 NHC). The results shown here are from the meta-analysis. Results for individual datasets are shown in Extended Data 1-MWAS. Species that achieved FDR<0.05 and PHet>0.1 in meta-analysis were considered significant. (c) Conditional MWAS testing [PD vs NHC] and [constipated vs. not constipated in past 3 months] in one model. The test was conducted in each dataset and then meta-analyzed (N = DOD: 465 PD, 223 NHC; NGRC: 202 PD, 133 NHC; UAB: 298 PD, 174 NHC). Results are for PD-species associations that remained significant after adjusting for constipation in the meta-analysis. Results for individual datasets are shown in Extended Data 3-Constipation. Note that associations that lost significance are towards the bottom of the table, which is sorted by significance. Thus, loss of signal could also be due to insufficient power for the extra variable in the model. (d) Conditional MWAS testing [PD vs NHC] and [men vs. women] simultaneously, tested in each data set, then meta-analyzed (N = DOD: 487 PD, 232 NHC; NGRC: 210 PD, 134 NHC; UAB: 309 PD, 178 NHC). Results shown are PD-species associations that retained significance after adjusting for sex in meta-analysis. Results for individual datasets are shown in Extended Data 4-Sex. (e) Network analysis. To assess interdependence among species, pairwise correlation in the absolute abundance of species was calculated using SparCC. Species that were correlated at |r|>0.2 and P<0.05 were assigned to a cluster by the Louvain algorithm and numbered by the algorithm 1-26. Edges denote the number of species with which a given species is correlated. For example, Blautia wexlerae is in the polymicrobial cluster #9, and the change in its relative abundance correlates with 25 other species, see Figure 1. Networks. testing differential relative abundances of species in PD vs. NHC. Full results in Extended Data 1-MWAS. DOD, NGRC, UAB: the 3 datasets. FDR: Benjamini–Hochberg false FC: Fold change in relative abundance comparing PD to NHC. Empty cells: Result did not reach significance. Datasets: Meta-analysis required species to have been tested in at least two datasets. The majority were tested in all three datasets, as shown. P_Het: Test of heterogeneity across datasets. If <0.1 results are deemed unreliable as true representatives of underlying datasets. Cluster: Algorithmically defined groups of interconnected species whose abundance counts were correlated. Edge: Number of species that correlate with the said species.

| Species | a. Replication |  |  |  | Species | b. Meta-analysis |  |  |  | c. Adjusted for constipation |  | d. Adjusted for sex |  | e. Network analysis |  |
| --- | --- | --- | --- | --- | --- | --- | --- | --- | --- | --- | --- | --- | --- | --- | --- |
|  | DOD |  | NGRC |  |  | Datasets | P_Het | FDR | FC | FDR | FC | FDR | FC | Cluster | Edges |
|  | FDR | FC | FDR | FC |  |  |  |  |  |  |  |  |  |  |  |
| *Bifidobacterium_dentium* | 9E-07 | 4.06 | 1E-05 | 4.36 | *Bifidobacterium_dentium* | DOD,UAB,NGRC | 0.73 | 5E-18 | 3.89 | 2E-14 | 3.73 | 1E-16 | 3.89 | 14 | 13 |
| *Streptococcus_mutans* | 1E-06 | 3.09 | 1E-04 | 2.33 | *Blautia_wexlerae* | DOD,UAB,NGRC | 0.87 | 1E-16 | 0.24 | 1E-11 | 0.27 | 3E-15 | 0.23 | 9 | 25 |
| *Clostridium_leptum* | 1E-06 | 3.51 | 5E-02 | 2.76 | *Roseburia_intestinalis* | DOD,UAB,NGRC | 0.71 | 1E-16 | 0.17 | 3E-11 | 0.20 | 1E-17 | 0.14 | 9 | 26 |
| *Ruthenibacterium_lactatiformans* | 5E-06 | 2.13 | 9E-04 | 3.08 | *Streptococcus_mutans* | DOD,UAB,NGRC | 0.28 | 2E-13 | 2.41 | 5E-12 | 2.27 | 1E-09 | 2.32 | 14 | 13 |
| *Blautia_wexlerae* | 2E-05 | 0.26 | 2E-05 | 0.22 | *Anaerostipes_hadrus* | DOD,UAB,NGRC | 0.97 | 9E-08 | 0.35 | 7E-04 | 0.45 | 1E-09 | 0.29 | 9 | 41 |
| *Ruminococcaceae_bacterium_D5* | 6E-05 | 2.34 | 2E-01 | 1.82 | *Eisenbergiella_tayi* | DOD,UAB,NGRC | 0.89 | 9E-08 | 2.59 | 9E-04 | 1.96 | 2E-09 | 2.97 | 4 | 10 |
| *Roseburia_intestinalis* | 1E-04 | 0.21 | 1E-04 | 0.15 | *Fusicatenibacter_saccharivorans* | DOD,UAB,NGRC | 0.83 | 1E-06 | 0.32 | 7E-04 | 0.40 | 3E-08 | 0.27 | 9 | 42 |
| *Eisenbergiella_tayi* | 3E-04 | 2.65 | 1E-01 | 2.22 | *Ruthenibacterium_lactatiformans* | DOD,UAB,NGRC | 0.17 | 1E-06 | 2.14 | 1E-04 | 1.66 | 1E-09 | 2.51 | 4 | 13 |
| *Coprobacillus_cateniformis* | 6E-04 | 1.87 | 5E-03 | 2.81 | *Ruminococcaceae_bacterium_D5* | DOD,UAB,NGRC | 0.52 | 1E-06 | 2.04 | 8E-04 | 1.76 | 2E-05 | 1.93 | 9 | 20 |
| *Ruminococcus_lactaris* | 2E-03 | 0.43 | 4E-02 | 0.41 | *Eubacterium_rectale* | DOD,UAB,NGRC | 0.64 | 2E-06 | 0.30 | 7E-04 | 0.37 | 4E-10 | 0.21 | 9 | 26 |
| *Lactobacillus_salivarius* | 2E-03 | 1.88 | 2E-01 | 1.40 | *Methanobrevibacter_smithii* | DOD,UAB,NGRC | 0.36 | 2E-06 | 2.53 | 1E-03 | 2.05 | 5E-04 | 2.02 | 9 | 25 |
| *Collinsella_massiliensis* | 2E-03 | 1.87 | 2E-01 | 1.25 | *Cloacibacillus_evryensis* | DOD,UAB,NGRC | 0.70 | 2E-06 | 1.81 | 9E-04 | 1.61 | 3E-05 | 1.73 | 9 | 9 |
| *Clostridium_sp_CAG_299* | 3E-03 | 0.47 | 2E-01 | 0.66 | *Hungatella_hathewayi* | DOD,UAB,NGRC | 0.29 | 2E-06 | 2.43 | 1E-03 | 2.01 | 6E-09 | 3.03 | 4 | 34 |
| *Anaerostipes_hadrus* | 3E-03 | 0.33 | 1E-02 | 0.35 | *Alistipes_indistinctus* | DOD,UAB,NGRC | 0.52 | 6E-06 | 2.27 | 0.02 | 1.93 | 9E-04 | 1.89 | 9 | 20 |
| *Lactobacillus_fermentum* | 4E-03 | 2.19 | 1E-02 | 1.77 | *Roseburia_faecis* | DOD,UAB,NGRC | 0.59 | 1E-05 | 0.34 | 0.01 | 0.44 | 5E-07 | 0.29 | 9 | 27 |
| *Ruminococcus_callidus* | 4E-03 | 0.81 | 9E-02 | 0.75 | *Turicibacter_sanguinis* | DOD,UAB,NGRC | 0.46 | 1E-05 | 1.80 | 8E-05 | 1.82 | 2E-04 | 1.72 | 10 | 3 |
| *Megasphaera_sp_DISK_18* | 4E-03 | 1.68 | 1E-01 | 1.40 | *Actinomyces_sp_HPA0247* | DOD,UAB | 0.54 | 2E-05 | 1.66 | 8E-04 | 1.55 | 4E-05 | 1.73 | 14 | 6 |
| *Porphyromonas_asaccharolytica* | 5E-03 | 1.73 | 8E-03 | 3.77 | *Parabacteroides_distasonis* | DOD,UAB,NGRC | 0.26 | 2E-05 | 2.90 | 1E-03 | 2.36 | 8E-04 | 2.42 | 2 | 6 |
| *Fusicatenibacter_saccharivorans* | 5E-03 | 0.28 | 1E-02 | 0.30 | *Clostridium_innocuum* | DOD,UAB,NGRC | 0.72 | 2E-05 | 2.14 | 0.02 | 1.65 | 9E-09 | 2.81 | 4 | 28 |
| *Lactobacillus_paragasseri* | 5E-03 | 1.59 | 2E-02 | 1.67 | *Bifidobacterium_longum* | DOD,UAB,NGRC | 0.48 | 3E-05 | 2.75 | 6E-04 | 2.87 | 2E-05 | 2.93 | 18 | 5 |
| *Faecalibacterium_prausnitzii* | 6E-03 | 0.40 | 1E-01 | 0.44 | *Faecalibacterium_prausnitzii* | DOD,UAB,NGRC | 0.87 | 7E-05 | 0.44 | 0.01 | 0.56 | 2E-07 | 0.34 | 9 | 43 |
| *Methanobrevibacter_smithii* | 7E-03 | 2.71 | 7E-03 | 3.63 | *Pseudoflavonifractor_sp_An184* | DOD,UAB,NGRC | 0.28 | 9E-05 | 1.60 | 0.01 | 1.45 | 3E-04 | 1.58 | 5 | 4 |
| *Hungatella_hathewayi* | 7E-03 | 2.21 | 5E-03 | 4.00 | *Ruminococcus_bicirculans* | DOD,UAB,NGRC | 0.58 | 9E-05 | 0.41 | 0.01 | 0.48 | 5E-06 | 0.35 | 9 | 27 |
| *Cloacibacillus_evryensis* | 7E-03 | 1.69 | 9E-03 | 2.09 | *Eubacterium_callanderi* | DOD,UAB,NGRC | 0.39 | 1E-04 | 1.31 | 0.02 | 1.21 | 8E-05 | 1.35 | 9 | 1 |
| *Alistipes_indistinctus* | 7E-03 | 2.52 | 1E-02 | 2.78 | *Lactobacillus_rhamnosus* | DOD,UAB,NGRC | 0.66 | 1E-04 | 1.53 | 1E-03 | 1.50 | 4E-05 | 1.60 | 18 | 8 |
| *Acidaminococcus_intestini* | 1E-02 | 2.98 | 2E-01 | 1.83 | *Eubacterium_limosum* | DOD,UAB,NGRC | 0.28 | 1E-04 | 1.43 |  |  | 4E-06 | 1.51 | none | 0 |
| *Lactobacillus_rhamnosus* | 1E-02 | 1.71 | 6E-02 | 1.51 | *Bifidobacterium_adolescentis* | DOD,UAB,NGRC | 0.25 | 2E-04 | 2.60 | 0.01 | 2.51 | 3E-03 | 2.14 | 18 | 3 |
| *Bifidobacterium_pullorum* | 2E-02 | 1.49 | 4E-02 | 1.30 | *Corynebacterium_amycolatum* | UAB,NGRC | 0.80 | 2E-04 | 2.34 | 2E-03 | 2.28 | 0.02 | 1.74 | 25 | 14 |
| *Lactobacillus_gasseri* | 2E-02 | 1.33 | 2E-03 | 2.02 | *Ruminococcus_callidus* | DOD,NGRC | 0.59 | 2E-04 | 0.80 | 7E-04 | 0.79 | 3E-04 | 0.79 | none | 0 |
| *Eubacterium_rectale* | 2E-02 | 0.38 | 4E-02 | 0.29 | *Eubacterium_sp_CAG_38* | DOD,UAB,NGRC | 0.57 | 2E-04 | 0.50 | 0.01 | 0.55 | 2E-04 | 0.48 | 9 | 18 |
| *Roseburia_faecis* | 3E-02 | 0.37 | 2E-02 | 0.22 | *Lawsonella_clevelandensis* | UAB,NGRC | 0.53 | 2E-04 | 1.63 | 7E-04 | 1.69 | 4E-03 | 1.49 | 25 | 34 |
| *Ruminococcus_bicirculans* | 3E-02 | 0.42 | 2E-02 | 0.30 | *Ruminococcus_lactaris* | DOD,UAB,NGRC | 0.28 | 3E-04 | 0.50 | 0.02 | 0.59 | 8E-06 | 0.47 | 9 | 16 |
| *Rothia_mucilaginosa* | 3E-02 | 0.59 | 5E-02 | 0.69 | *Bifidobacterium_bifidum* | DOD,UAB,NGRC | 0.40 | 3E-04 | 1.98 | 2E-03 | 1.86 | 9E-04 | 1.92 | 18 | 3 |
| *Christensenella_minuta* | 3E-02 | 1.47 | 8E-02 | 1.29 | *Roseburia_inulinivorans* | DOD,UAB,NGRC | 0.44 | 3E-04 | 0.43 | 4E-03 | 0.47 | 6E-07 | 0.31 | 9 | 31 |
| *Enorma_massiliensis* | 3E-02 | 1.61 | 2E-01 | 1.27 | *Eubacterium_ramulus* | DOD,UAB,NGRC | 0.74 | 4E-04 | 0.57 | 0.01 | 0.63 | 1E-05 | 0.50 | 9 | 22 |
| *Bifidobacterium_bifidum* | 3E-02 | 2.44 | 5E-02 | 2.53 | *Christensenella_minuta* | DOD,UAB,NGRC | 0.71 | 4E-04 | 1.33 | 0.02 | 1.26 | 3E-04 | 1.35 | 4 | 1 |
| *Parabacteroides_distasonis* | 3E-02 | 2.18 | 3E-04 | 4.36 | *Collinsella_stercoris* | DOD,UAB,NGRC | 0.76 | 4E-04 | 1.79 | 3E-03 | 1.74 | 0.01 | 1.60 | 14 | 10 |
| *Pseudoflavonifractor_sp_An184* | 3E-02 | 1.49 | 2E-02 | 2.35 | *Bifidobacterium_pullorum* | DOD,NGRC | 0.41 | 4E-04 | 1.37 | 0.01 | 1.29 | 8E-04 | 1.36 | 18 | 10 |
| *Roseburia_inulinivorans* | 3E-02 | 0.45 | 3E-02 | 0.28 | *Lactobacillus_fermentum* | DOD,UAB,NGRC | 0.22 | 4E-04 | 1.71 | 7E-04 | 1.63 | 3E-04 | 1.61 | 14 | 13 |
| *Clostridium_innocuum* | 3E-02 | 1.88 | 4E-02 | 2.62 | *Bifidobacterium_breve* | DOD,UAB,NGRC | 0.17 | 5E-04 | 1.66 | 4E-05 | 1.66 | 2E-05 | 1.62 | 18 | 16 |
| *Alistipes_finegoldii* | 4E-02 | 2.14 | 1E-01 | 2.68 | *Agathobaculum_butyriciproducens* | DOD,UAB,NGRC | 0.42 | 6E-04 | 0.50 | 0.04 | 0.61 | 5E-06 | 0.39 | 9 | 35 |
| *Eubacterium_ramulus* | 5E-02 | 0.53 | 6E-02 | 0.51 | *Veillonella_dispar* | DOD,UAB,NGRC | 0.82 | 6E-04 | 0.81 | 0.03 | 0.85 | 2E-04 | 0.79 | 6 | 5 |
| *Catabacter_hongkongensis* | 5E-02 | 1.36 | 2E-01 | 1.24 | *Acidaminococcus_intestini* | DOD,UAB,NGRC | 0.47 | 7E-04 | 2.11 | 3E-03 | 2.11 | 0.03 | 1.66 | 14 | 2 |
|  |  |  |  |  | *Megasphaera_sp_DISK_18* | DOD,UAB,NGRC | 0.27 | 7E-04 | 1.41 | 5E-03 | 1.33 | 0.01 | 1.34 | 8 | 1 |
|  |  |  |  |  | *Bifidobacterium_pseudocatenulatum* | DOD,UAB,NGRC | 0.65 | 1E-03 | 1.78 | 0.01 | 1.70 | 1E-03 | 1.79 | 18 | 2 |
|  |  |  |  |  | *Lactobacillus_paragasseri* | DOD,UAB,NGRC | 0.20 | 1E-03 | 1.46 | 8E-04 | 1.39 | 9E-04 | 1.44 | 14 | 9 |
|  |  |  |  |  | *Peptoniphilus_sp_HMSC062D09* | UAB,NGRC | 1.00 | 1E-03 | 1.91 | 1E-03 | 2.04 | 0.01 | 1.73 | 25 | 29 |
|  |  |  |  |  | *Streptococcus_anginosus_group* | DOD,UAB,NGRC | 0.71 | 1E-03 | 1.40 | 0.04 | 1.28 | 9E-04 | 1.42 | 14 | 11 |
|  |  |  |  |  | *Rothia_mucilaginosa* | DOD,UAB,NGRC | 0.45 | 1E-03 | 0.69 | 0.03 | 0.75 | 5E-04 | 0.66 | 14 | 13 |
|  |  |  |  |  | *Alistipes_finegoldii* | DOD,UAB,NGRC | 0.79 | 1E-03 | 2.13 |  |  | 2E-03 | 2.16 | 9 | 17 |
|  |  |  |  |  | *Anaerococcus_vaginalis* | UAB,NGRC | 0.77 | 1E-03 | 1.78 | 0.01 | 1.78 | 0.01 | 1.67 | 25 | 34 |
|  |  |  |  |  | *Scardovia_wiggisiae* | DOD,UAB | 0.25 | 1E-03 | 1.36 | 0.01 | 1.31 | 4E-03 | 1.35 | 14 | 8 |
|  |  |  |  |  | *Prevotella_buccalis* | UAB,NGRC | 0.22 | 1E-03 | 2.31 | 0.01 | 2.34 | 0.03 | 2.37 | 25 | 26 |
|  |  |  |  |  | *Catabacter_hongkongensis* | DOD,UAB,NGRC | 0.85 | 1E-03 | 1.29 | 0.02 | 1.24 | 3E-04 | 1.35 | 4 | 2 |
|  |  |  |  |  | *Porphyromonas_asaccharolytica* | DOD,UAB,NGRC | 0.13 | 2E-03 | 1.99 | 0.02 | 1.72 | 0.02 | 1.92 | 25 | 28 |
|  |  |  |  |  | *Campylobacter_ureolyticus* | UAB,NGRC | 0.38 | 2E-03 | 1.54 |  |  | 0.01 | 1.90 | 25 | 29 |
|  |  |  |  |  | *Atopobium_minutum* | UAB,NGRC | 0.67 | 2E-03 | 1.38 | 0.01 | 1.39 | 9E-04 | 1.43 | 25 | 32 |
|  |  |  |  |  | *Lactobacillus_salivarius* | DOD,UAB,NGRC | 0.20 | 2E-03 | 1.49 | 0.01 | 1.41 | 9E-04 | 1.52 | 14 | 6 |
|  |  |  |  |  | *Eubacterium_hallii* | DOD,UAB,NGRC | 0.17 | 2E-03 | 0.44 | 0.02 | 0.55 | 7E-06 | 0.35 | 9 | 46 |
|  |  |  |  |  | *Monoglobus_pectinilyticus* | DOD,UAB,NGRC | 0.87 | 2E-03 | 0.55 | 0.01 | 0.58 | 0.01 | 0.58 | 7 | 1 |
|  |  |  |  |  | *Veillonella_infantium* | DOD,UAB,NGRC | 0.34 | 3E-03 | 0.82 |  |  | 4E-03 | 0.81 | 6 | 5 |
|  |  |  |  |  | *Bacteroides_coagulans* | UAB,NGRC | 0.76 | 3E-03 | 1.57 | 0.01 | 1.56 | 1E-03 | 1.68 | 25 | 29 |
|  |  |  |  |  | *Harryflintia_acetispora* | DOD,UAB,NGRC | 0.87 | 4E-03 | 1.42 | 0.02 | 1.38 | 0.01 | 1.39 | 9 | 5 |
|  |  |  |  |  | *Butyricimonas_virosa* | DOD,UAB,NGRC | 0.75 | 4E-03 | 1.63 |  |  |  |  | 9 | 15 |
|  |  |  |  |  | *Porphyromonas_sp_HMSC065F10* | UAB,NGRC | 0.73 | 4E-03 | 1.95 | 0.01 | 2.07 | 4E-03 | 2.04 | 25 | 29 |
|  |  |  |  |  | *Gordonibacter_pamelaeae* | DOD,UAB,NGRC | 0.75 | 5E-03 | 1.54 | 0.03 | 1.45 | 4E-05 | 1.90 | 4 | 35 |
|  |  |  |  |  | *Bacteroides_caccae* | DOD,UAB,NGRC | 0.47 | 5E-03 | 1.90 |  |  | 0.04 | 1.64 | 2 | 5 |
|  |  |  |  |  | *Roseburia_sp_CAG_309* | DOD,UAB,NGRC | 0.69 | 0.01 | 0.82 | 0.03 | 0.83 | 8E-04 | 0.78 | none | 0 |
|  |  |  |  |  | *Faecalitalea_cylindroides* | DOD,UAB,NGRC | 0.93 | 0.01 | 1.33 | 0.05 | 1.27 | 0.04 | 1.26 | none | 0 |
|  |  |  |  |  | *Clostridium_sp_CAG_58* | DOD,UAB,NGRC | 0.83 | 0.01 | 0.58 | 0.05 | 0.62 | 5E-03 | 0.55 | 9 | 27 |
|  |  |  |  |  | *Lawsonibacter_asaccharolyticus* | DOD,UAB,NGRC | 0.32 | 0.01 | 0.61 |  |  | 0.01 | 0.62 | 9 | 24 |
|  |  |  |  |  | *Anaerotruncus_colihominis* | DOD,UAB,NGRC | 0.57 | 0.01 | 1.65 |  |  | 6E-06 | 2.36 | 4 | 22 |
|  |  |  |  |  | *Actinomyces_turicensis* | UAB,NGRC | 0.61 | 0.01 | 1.39 | 0.03 | 1.36 | 4E-03 | 1.45 | 25 | 24 |
|  |  |  |  |  | *Bilophila_wadsworthia* | DOD,UAB,NGRC | 0.75 | 0.01 | 1.64 |  |  |  |  | 9 | 3 |
|  |  |  |  |  | *Eubacterium_sp_CAG_274* | DOD,UAB,NGRC | 0.66 | 0.01 | 0.80 |  |  | 5E-03 | 0.77 | none | 0 |
|  |  |  |  |  | *Collinsella_massiliensis* | DOD,UAB,NGRC | 0.10 | 0.01 | 1.42 | 0.01 | 1.31 | 0.02 | 1.29 | 14 | 6 |
|  |  |  |  |  | *Pseudomonas_fragi* | UAB,NGRC | 0.80 | 0.02 | 0.69 | 0.05 | 0.71 | 0.02 | 0.69 | 21 | 1 |
|  |  |  |  |  | *Varibaculum_cambriense* | UAB,NGRC | 0.61 | 0.02 | 1.50 | 0.02 | 1.57 | 5E-03 | 1.62 | 25 | 34 |
|  |  |  |  |  | *Cloacibacillus_porcorum* | DOD,UAB,NGRC | 0.23 | 0.02 | 1.25 |  |  | 0.01 | 1.22 | none | 0 |
|  |  |  |  |  | *Haemophilus_parainfluenzae* | DOD,UAB,NGRC | 0.29 | 0.02 | 0.85 |  |  | 0.01 | 0.82 | 6 | 6 |
|  |  |  |  |  | *Eubacterium_eligens* | DOD,UAB,NGRC | 0.83 | 0.02 | 0.61 |  |  | 4E-04 | 0.47 | 9 | 51 |
|  |  |  |  |  | *Lachnoclostridium_sp_An138* | DOD,UAB,NGRC | 1.00 | 0.02 | 1.25 |  |  | 0.01 | 1.30 | 5 | 4 |
|  |  |  |  |  | *Peptococcus_niger* | UAB,NGRC | 0.72 | 0.02 | 1.27 |  |  | 0.02 | 1.29 | 25 | 24 |
|  |  |  |  |  | *Massiliomicrobiota_timonensis* | UAB,NGRC | 0.71 | 0.03 | 1.18 |  |  | 0.02 | 1.20 | none | 0 |
|  |  |  |  |  | *Streptococcus_lutetiensis* | DOD,UAB | 0.38 | 0.03 | 1.24 |  |  |  |  | 18 | 2 |
|  |  |  |  |  | *Veillonella_atypica* | DOD,UAB,NGRC | 0.99 | 0.03 | 0.82 |  |  | 0.02 | 0.81 | 6 | 6 |
|  |  |  |  |  | *Clostridium_sp_CAG_253* | DOD,UAB,NGRC | 0.73 | 0.03 | 0.86 |  |  | 0.04 | 0.86 | none | 0 |
|  |  |  |  |  | *Clostridium_scindens* | DOD,UAB,NGRC | 0.96 | 0.03 | 1.34 |  |  | 2E-04 | 1.66 | 4 | 43 |
|  |  |  |  |  | *Candidatus_Methanomassiliicoccus_inte* | DOD,NGRC | 0.34 | 0.03 | 1.26 |  |  |  |  | 9 | 3 |
|  |  |  |  |  | *Lachnospira_pectinoschiza* | DOD,UAB,NGRC | 0.13 | 0.03 | 0.52 |  |  | 0.03 | 0.37 | 9 | 20 |
|  |  |  |  |  | *Finegoldia_magna* | UAB,NGRC | 0.71 | 0.03 | 1.74 | 0.02 | 1.97 |  |  | 25 | 37 |
|  |  |  |  |  | *Porphyromonas_uenonis* | UAB,NGRC | 0.32 | 0.03 | 1.28 |  |  |  |  | 25 | 30 |
|  |  |  |  |  | *Dorea_longicatena* | DOD,UAB,NGRC | 0.86 | 0.04 | 0.65 |  |  | 4E-04 | 0.49 | 9 | 47 |
|  |  |  |  |  | *Roseburia_hominis* | DOD,UAB,NGRC | 0.50 | 0.04 | 0.63 |  |  | 0.01 | 0.56 | 9 | 34 |
|  |  |  |  |  | *Peptoniphilus_lacrimalis* | UAB,NGRC | 0.89 | 0.04 | 1.49 |  |  |  |  | 25 | 35 |
|  |  |  |  |  | *Akkermansia_muciniphila* | DOD,UAB,NGRC | 0.20 | 0.05 | 2.05 |  |  | 0.02 | 2.01 | 9 | 1 |

### Meta-analysis

We conducted meta-analyses of the MWAS results and identified 96 species, 1,119 gene families, and 95 pathways associated with PD (**Extended Data 2-MWAS**). We used random-effects models and set significance as FDR < 0.05 with no evidence of heterogeneity (P_Het_ > 0.1). Henceforth, when referring to an association, we use the results of meta-analysis. Of the 96 species that had significantly altered abundances in PD vs. NHC, 65 were elevated in PD and 31 were reduced (**Table 2. MWAS**). Similarly, of the 1,119 altered gene families, 690 were elevated, and 429 were reduced in PD, and of the 95 altered pathways, 60 were elevated and 35 reduced in PD. These results indicate a widespread dysbiosis involving ∼35% of species and ∼20% of genes and pathways, and a predominance of species enrichment over depletion in PD. However, while enriched species outnumber depleted ones, i.e., there is more diversity in the species that are enriched than those that are depleted, gain is a less prevalent dysbiotic feature (seen in 10-30% of patients) than loss (detected in 70% of patients) (see **Microbial features as biomarkers for subtyping** section). Not surprisingly, the taxa that are most easily detected across published literature had the strongest signals, namely *Bifidobacterium dentium,* which was elevated in PD by 4-fold (FDR=5E-18), and *Roseburia intestinalis*, which was reduced in PD by 6-fold (FDR=1E-16). We confirmed most of the findings of the recent meta-analysis ^15^ and expanded on it by ∼50 additional PD-associated species, including 16 rare opportunistic pathogens that form an interconnected polymicrobial cluster (**Table 2. MWAS**).

A limitation of this study was the inability to accurately estimate the relative abundance of *E. coli* in NGRC and UAB samples. *E. coli* is elevated in the PD gut, as shown in DOD datasets and others ^15,22^. Alerted by a strong heterogeneity signal and the lack of association with PD in NGRC and UAB, we noted that the relative abundance of *E. coli* had increased to >15% in NGRC and UAB, compared to ∼1% in DOD (**Extended Data 2-MWAS**). We suspect that the swab stool collection kit used for NGRC and UAB allowed *E. coli* to grow uncontrollably during shipping from subjects’ homes to the lab. *Klebsiella* spp, which are related to *E. coli*, had also bloomed but to a lesser degree. Other *E. coli*-related species, e.g., *Shigella, Salmonella*, and *Citrobacter*, were either too rare to detect and test, or, if tested, did not show a PD signal. Observations made about *E. coli* and *Klebsiella* spp will therefore be limited to the DOD dataset.

### Constipation is not the sole cause of PD dysbiosis

Constipation has a large and reproducible effect on the composition of the gut microbiome ^16,23^. Constipation is also an early and common feature of PD that persists through disease progression ^3^. In this study, constipation was 4-16 times more prevalent in patients than in controls (**Table 1. Participant characteristics**). To investigate the effect of constipation on the composition of microbiome, we conducted MWAS in PD and in NHC separately, testing the association of constipation history in the past 3 months with the relative abundance of species (**Extended Data 3-Constipation**). Constipation affected relative abundances of 16 species in PD and only one species in NHC (**Table 3. Constipation & Sex**). To determine if the differential abundance of species in PD vs. NHC was due to constipation, we ran a conditional MWAS where we tested the association of species with PD status and with constipation simultaneously, estimating the effect of each variable (PD and constipation) on the species relative abundance while controlling for the other variable (**Extended Data 3-Constipation**). If a PD-microbe association is secondary to constipation, the signal for association with PD will fade when constipation is in the model. Analyses were conducted in each dataset and then meta-analyzed. 69 of 96 PD-associated species remained significantly associated with PD at FDR < 0.05 (**Table 2. MWAS**). *Akkermansia muciniphila* and *Alistipes finegoldii* lost significance. *Eisenbergiella tayi* and *Faecalibacterium prausnitzii,* which had the strongest association with constipation in PD, retained their significance for association with PD in addition to their association with constipation. In sum, constipation contributes to dysbiosis in PD, yet 72% of the dysbiotic features have a robust signal for PD independent of constipation.

**Table 3.** Constipation and Sex. Testing the effects of constipation and sex on the microbiome. MWAS was conducted in PD and neurologically healthy controls (NHC) separately, once to test the effect of constipation and once to test the effect of sex on the relative abundance of species. Constipation was defined as having <3 bowel movements per week in the past three months (yes/no). Shown are significant associations (FDR<0.05, P Het>0.1). For example, among individuals with PD, those who reported constipation had 2.62-fold higher abundance of Eisenbergiella tayi and 2.68-fold lower abundance of Faecalibacterium prausnitzii than those without constipation. Full results are in Extended Data 3-Constipation, and Extended Data 4-Sex.

| Species | FDR | FC | Species | FDR | FC |
| --- | --- | --- | --- | --- | --- |
| **a. Constipation in PD metagenomes (N PD = 465 DOD, 202 NGRC, 298 UAB)** |  |  |  |  |  |
| *Eisenbergiella tayi* | 2E-04 | 2.62 | *Actinomyces sp ICM47* | 4E-03 | 0.64 |
| *Clostridium innocuum* | 1E-03 | 2.27 | *Veillonella parvula* | 5E-03 | 0.60 |
| *Ruthenibacterium lactatiformans* | 2E-05 | 2.09 | *Ruminococcus lactaris* | 3E-03 | 0.53 |
| *Bacteroides massiliensis* | 0.04 | 1.92 | *Agathobaculum butyriciproducens* | 0.01 | 0.47 |
| *Hungatella hathewayi* | 0.03 | 1.85 | *Fusicatenibacter saccharivorans* | 0.02 | 0.44 |
| *Clostridium leptum* | 0.02 | 1.82 | *Roseburia faecis* | 0.02 | 0.43 |
| *Eggerthella lenta* | 0.02 | 1.82 | *Anaerostipes hadrus* | 9E-04 | 0.39 |
| *Clostridium hylemonae* | 0.02 | 1.22 | *Faecalibacterium prausnitzii* | 3E-04 | 0.37 |
| **b. Constipation in NHC metagenomes (N NHC = 223 DOD, 133 NGRC, 174 UAB)** |  |  |  |  |  |
| Ruminococcaceae bacterium D5 | 0.01 | 2.42 |  |  |  |
| **c. Sex in PD metagenomes (N PD = 487 DOD, 210 NGRC, 309 UAB)** |  |  |  |  |  |
| *Oscillibacter sp 57 20* | 3E-08 | 5.88 | *Bacteroides finegoldii* | 0.02 | 1.67 |
| *Oscillibacter sp CAG 241* | 2E-06 | 4.09 | *Parabacteroides goldsteinii* | 0.03 | 1.61 |
| *Parabacteroides merdae* | 1E-05 | 4.08 | *Holdemanella biformis* | 7E-05 | 1.61 |
| *Firmicutes bacterium CAG 83* | 2E-06 | 4.04 | *Ruminococcaceae bacterium D5* | 0.03 | 1.58 |
| *Barnesiella intestinihominis* | 2E-06 | 3.89 | *Eubacterium ramulus* | 0.05 | 1.50 |
| *Collinsella aerofaciens* | 2E-04 | 3.43 | *Bacteroides faecis* | 0.03 | 1.49 |
| *Blautia obeum* | 2E-06 | 3.41 | *Dialister sp CAG 357* | 0.01 | 1.48 |
| *Bacteroides stercoris* | 1E-04 | 3.21 | *Collinsella massiliensis* | 0.01 | 1.39 |
| *Corynebacterium amycolatum* | 4E-04 | 3.20 | *Megasphaera sp DISK 18* | 0.02 | 1.35 |
| *Gemmiger formicilis* | 2E-06 | 3.19 | *Slackia isoﬂavoniconvertens* | 0.02 | 1.34 |
| *Alistipes shahii* | 1E-04 | 3.06 | *Catenibacterium mitsuokai* | 0.01 | 1.31 |
| *Dorea formicigenerans* | 4E-06 | 2.93 | *Phascolarctobacterium succinatutens* | 2E-04 | 1.31 |
| *Eubacterium siraeum* | 7E-04 | 2.90 | *Roseburia sp CAG 309* | 0.03 | 1.20 |
| *Faecalibacterium prausnitzii* | 7E-05 | 2.82 | *Anaerofustis stercorihominis* | 0.03 | 0.79 |
| *Methanobrevibacter smithii* | 2E-04 | 2.81 | *Clostridiales bacterium 1 7 47FAA* | 0.01 | 0.72 |
| *Eubacterium eligens* | 7E-05 | 2.75 | *Murimonas intestini* | 5E-03 | 0.72 |
| *Alistipes putredinis* | 1E-03 | 2.71 | *Ruthenibacterium lactatiformans* | 2E-03 | 0.64 |
| *Coprococcus catus* | 2E-03 | 2.49 | *Clostridium aldenense* | 7E-04 | 0.63 |
| *Eubacterium rectale* | 5E-03 | 2.49 | *Intestinibacter bartlettii* | 0.01 | 0.62 |
| *Roseburia inulinivorans* | 5E-04 | 2.42 | *Blautia coccoides* | 0.02 | 0.58 |
| *Alistipes indistinctus* | 2E-04 | 2.34 | *Clostridium bolteae* | 0.02 | 0.57 |
| *Ruminococcus torques* | 3E-03 | 2.26 | *Clostridium lavalense* | 0.01 | 0.56 |
| *Dorea longicatena* | 1E-03 | 2.24 | *Peptoniphilus harei* | 0.01 | 0.56 |
| *Agathobaculum butyriciproducens* | 2E-03 | 2.18 | *Hungatella hathewayi* | 0.01 | 0.52 |
| *Eubacterium hallii* | 3E-03 | 2.17 | *Eisenbergiella massiliensis* | 0.01 | 0.51 |
| *Bacteroides massiliensis* | 0.01 | 2.16 | *Clostridium citroniae* | 0.01 | 0.49 |
| *Ruminococcus bicirculans* | 0.01 | 2.14 | *Blautia sp CAG 257* | 0.01 | 0.48 |
| *Butyricimonas virosa* | 0.03 | 2.13 | *Clostridium scindens* | 4E-06 | 0.45 |
| *Adlercreutzia equolifaciens* | 1E-03 | 2.13 | *Flavonifractor plautii* | 2E-06 | 0.43 |
| *Butyricimonas synergistica* | 2E-04 | 2.05 | *Ruminococcus gnavus* | 5E-03 | 0.43 |
| *Bilophila wadsworthia* | 2E-03 | 2.03 | *Erysipelatoclostridium ramosum* | 5E-06 | 0.42 |
| *Anaerostipes hadrus* | 0.03 | 1.91 | *Clostridium clostridioforme* | 4E-05 | 0.40 |
| *Eubacterium sp CAG 180* | 2E-03 | 1.91 | *Clostridium innocuum* | 4E-05 | 0.40 |
| *Asaccharobacter celatus* | 0.01 | 1.87 | *Clostridium symbiosum* | 3E-05 | 0.37 |
| *Paraprevotella xylaniphila* | 0.01 | 1.80 | *Eggerthella lenta* | 5E-05 | 0.26 |
| **d. Sex in NHC metagenomes** (N NHC = 232 DOD, 134 NGRC, 178 UAB) |  |  |  |  |  |
| *Eubacterium rectale* | 7E-04 | 4.33 | *Firmicutes bacterium CAG 145* | 0.01 | 0.42 |
| *Coprococcus comes* | 3E-03 | 2.84 | *Clostridium bolteae* | 0.02 | 0.42 |
| *Dorea longicatena* | 0.01 | 2.13 | *Clostridium symbiosum* | 0.01 | 0.41 |
| *Corynebacterium amycolatum* | 0.04 | 2.08 | *Sellimonas intestinalis* | 0.03 | 0.41 |
| *Dielma fastidiosa* | 0.03 | 0.67 | *Flavonifractor plautii* | 2E-03 | 0.41 |
| *Lachnoclostridium* sp An138 | 2E-03 | 0.63 | *Anaerotruncus colihominis* | 2E-03 | 0.35 |
| *Ruthenibacterium lactatiformans* | 0.01 | 0.53 | *Anaerotignum lactatifermentans* | 0.01 | 0.35 |
| *Gordonibacter pamelaee* | 0.01 | 0.48 | *Eisenbergiella massiliensis* | 7E-04 | 0.27 |
| *Eisenbergiella tayi* | 0.01 | 0.45 | *Intestinimonas butyriciproducens* | 1E-03 | 0.26 |

### Strong sex effect on PD microbiome

Our patient datasets are ∼60% men due to the higher prevalence of PD in men ^24^, and our controls are ∼60% women because most controls are spouses of patients. Given this structure, if a microbe were more prevalent in one sex, it could generate a false signal for the association of the microbe with disease. First, we conducted MWAS to test the association of sex with species abundance in PD and controls separately, in each of the three datasets, and then meta-analyzed (**Extended Data 4-Sex**). Sex affected differential abundances of 70 species in PD and only 18 species in NHC (**Table 3. Constipation & Sex**). Second, to assess potential confounding by sex on PD-species associations, we conducted conditional MWAS with both PD and sex in the model, in individual datasets, and then meta-analyzed (**Extended Data 4-Sex**). In sex-adjusted meta-analysis, 89 of 96 PD-associated species remained significantly associated with PD (**Table 2. MWAS**). In sum, data exhibited a greater sex effect on the microbiome of individuals with PD than those without PD, which suggests a synergistic play of sex and PD on the gut microbiome. Moreover, despite the sex effect in PD microbiome, 93% of PD-species associations were robust to confounding by sex.

### A bird’s-eye view of PD microbiome

Stepping back to the big picture, a total of 1,131 species were detected, 272 passed thresholds for inclusion in each dataset for analysis and then in meta-analysis, and 96 had altered abundances in PD. We were interested in seeing the overall picture, with interconnectivity of species depicted in a microbial network, and determining if and which species form polymicrobial clusters that grow in abundance together or are lost together (**Figure 1. Network**). To create the networks, treating PD and NHC separately, we calculated pairwise correlation in absolute counts of all 1,131 species using SparCC ^25^, setting a threshold for correlation at |r|>0.2 with P < 0.05. We then used the Louvain algorithm^26^ to assign species to polymicrobial clusters according to their correlations. A cluster is therefore a group of species whose abundances increase together or decrease together (positive correlation) or compete (negative correlation). We plotted the clusters to create the network in the PD and NHC metagenomes and colored the species that are elevated or reduced in the PD meta-analysis to highlight them in the network (**Extended Data 5-Networks)**.

**Figure 1.**
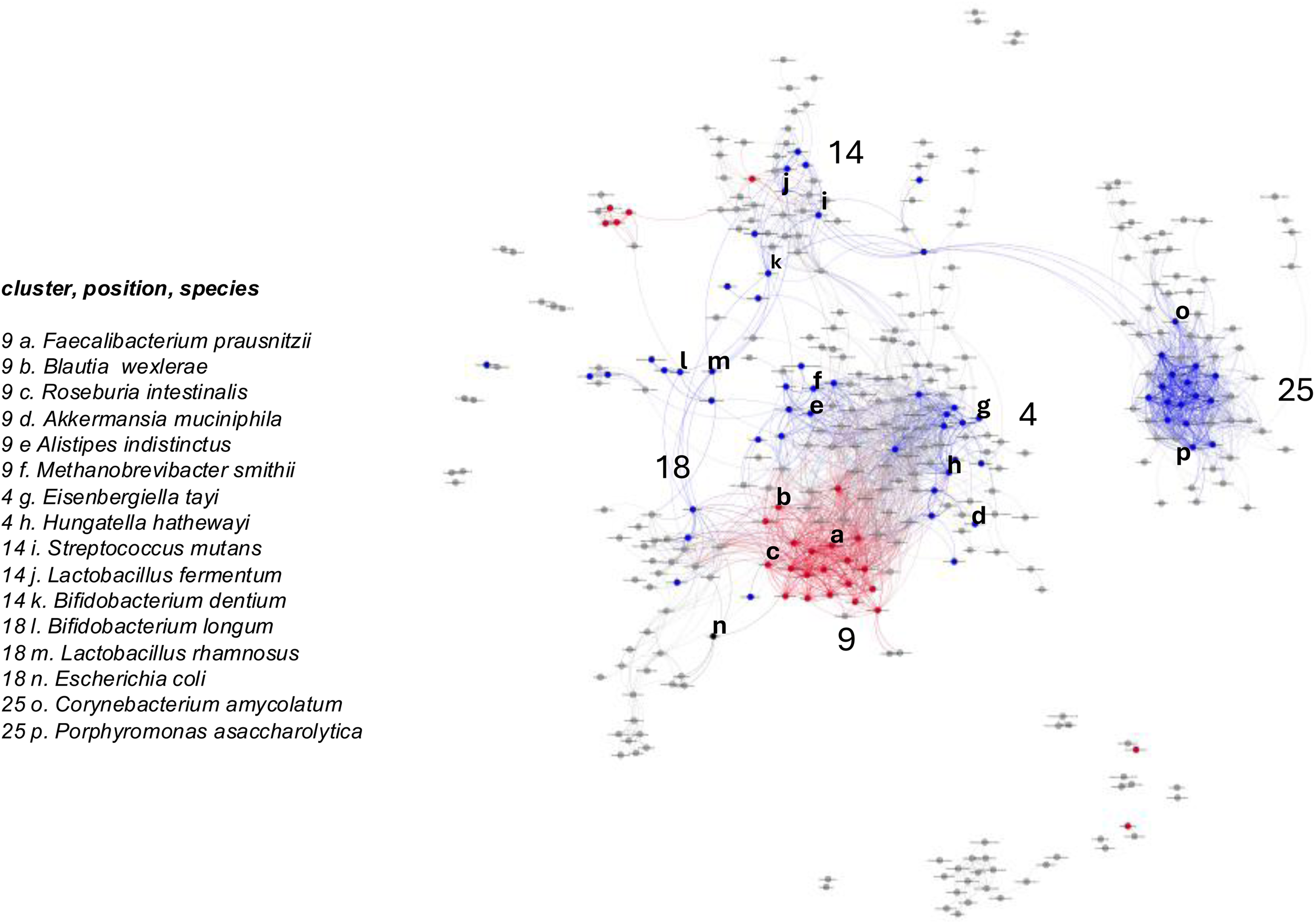
Network of polymicrobial clusters in PD gut. A bird’s eye view of PD gut microbiome. Species that were detected in the PD gut metagenome are plotted according to correlation in their abundance counts, creating a visual of correlated polymicrobial clusters. Circles are species, colored as red if they are reduced in PD, blue if they are elevated in PD, and gray if not associated with PD. Lines connect species whose abundances are correlated. For example, the intense central red is a polymicrobial cluster of mostly fiber degraders; the intense blue island is a polymicrobial cluster of opportunistic pathogens. Twenty-six clusters were called; a few are annotated by number and representative species. Full results, and side-by-side plots of PD and NHC networks are in **Extended Data 5-Networks**.

Twenty-six clusters were called in the PD microbiome (see **Figure 1. Network** for visual presentation, and **Table 2. MWAS** for cluster assignment of species that are associated with PD). The most striking clusters were 9 and 25; also of note were clusters 4, 14, and 18. Cluster 9 was composed of 63 species, 20 of which were depleted in PD, and 11 were elevated in PD. The depleted species are predominantly beneficial bacteria that degrade fiber and produce anti-inflammatory molecules and SCFAs, including *Eubacterium* spp, *Roseburia* spp, *Blautia wexlerae,* and *Faecalibacterium prausnitzii*. Notable species that are enriched in this cluster include *Akkermansia muciniphila* and *Methanobrevibacter smithii,* both of which reduce gut motility, and *Alistipes* spp, which can be both pathogenic and protective of the gut. In short, a drastic loss of bacteria that degrade fibers is correlated with an increase in an array of microbes with diverse effects. This observation suggests that if depletion of fiber degraders has caused the increase in the enriched species in cluster 9, a prebiotic or probiotic dietary supplementation to replenish fiber degraders ^27–29^ may also restore some of the abnormally abundant microbes, achieving wider benefit.

Cluster 25 is a large polymicrobial cluster of 64 species that form an island with only one connection to the rest of the network, via *Streptococcus anginosus* (**Figure 1. Network**). Species in this cluster are rare opportunistic pathogens. Despite their rarity, 16 of them were detected and tested individually, and all were significantly elevated in PD (**Table 2. MWAS**). Opportunistic pathogens are commensal to the human microbiome (gut, mouth, or vagina), but in immunocompromised individuals, can cause infections and systemic and persistent inflammation. Cluster 25 included species from the genera *Corynebacterium*, *Prevotella*, *Porphyromonas*, *Peptoniphilus*, *Anaerococcus*, *Finegoldia*, *Lawsonella*, *Bacteroides*, *Campylobacter*, *Varibaculum*, *Actinomyces*, *Atopobium*, and *Peptococcus*. To speculate, we wonder if this cluster is similar in structure and function to the multigenus consortium in the microbiome of supragingival dental plaques, where *Corynebacterium* forms a tree-like scaffolding that supports the accumulation of other bacteria ^30^.

Clusters 4, 14, and 18 include an array of species that are enriched in PD. Cluster 4 includes 9 PD-associated species, all elevated in PD. Among these are bacteria known to be pathogens (*Eisenbergiella tayi*, *Catabacter hongkongensis*), beneficial (*Clostridium leptum*, *Christensenella minuta*), or to have both beneficial and harmful effects (*Hungatella hathewayi*, *Clostridium scindens*). Clusters 14 and 18 are where *Bifidobacteria*, *Lactobacillus*, *Streptococcus,* and some of the *Actinomyces* species are interspersed.

### Functional inference

Differential abundances of microbial pathways (PTW) and gene families (KO: Kyoto Encyclopedia of Genes and Genomes (KEGG) Ontology) are used to infer the functional impact of dysbiosis. Here, we expand these sources to also include species that contributed to each KO and PTW (**Extended Data 6-Inference**). 1,119 gene families and 95 pathways were significantly altered in the PD metagenome. To make data on over a thousand significant outcomes more manageable for discussion, we grouped them first by inferred function taken from KEGG, and then in larger groups of inferred relevance to microbiome (housekeeping, metabolism, homeostasis) or to PD (pathogenicity, neuroactive signaling, toxicity). Selected examples are discussed below and plotted in **Figure 2. Genes & pathways**.

**Figure 2.**
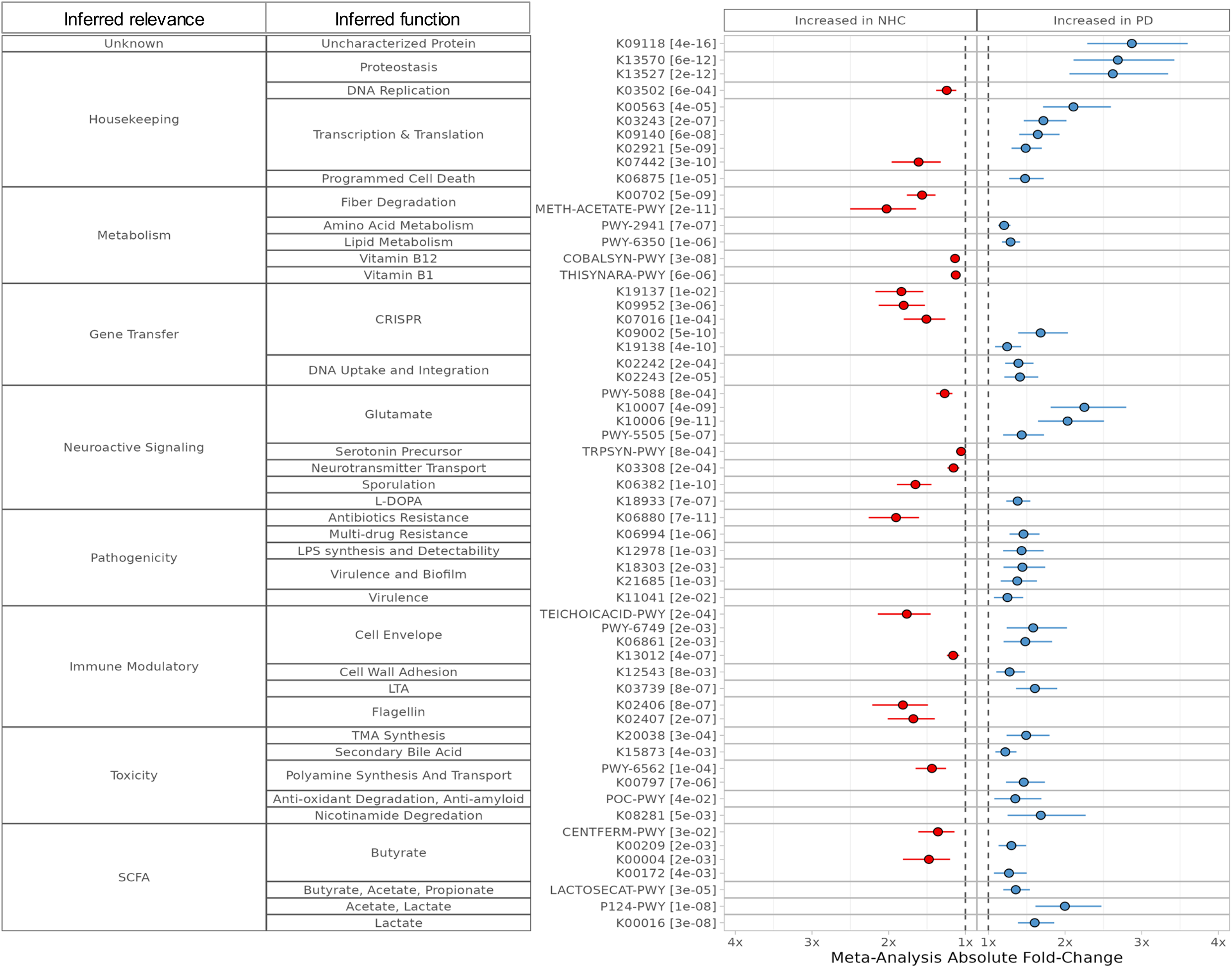
Genes & pathways. Widespread dysbiosis was detected in the PD microbiome, including altered abundances of 1119 microbial gene families (KO) and 95 pathways (PTW). Analysis included metagenome-wide association studies (MWAS) to test differential relative abundances of KO and PTW, in turn, in each of three datasets, followed by meta-analysis. Full results are in Extended Data 2-MWAS. We complemented the standard approach of using KO and PTW for functional inference by identifying the species that contributed to the loss or gain of each KO and PTW, see Extended Data 6-Function. Representative features are plotted here with their fold changes in PD vs. controls and standard errors, grouped by inferred function and relevance to PD pathobiology and microbiome homeostasis. Novel findings of note include significant imbalances in genes involved in the CRISPR system and DNA uptake, an increase in virulence, multidrug resistance, and a decline in immunomodulatory features in addition to confirming evidence for heightened pathogenicity, an increase in L-dopa breakdown by an archaea, and elevated synthesis of short-chain fatty acids including butyrate by *Bifidobacteria* spp, in contrast to the reduction in SCFA production due to depletion of fiber degrading bacteria.

If we rank the results by statistical significance, the most significant change was a 3-fold increase in an as-yet uncharacterized gene family (FDR=4E-16), which underscores that there is more to the PD microbiome than we currently understand. Nearly 500 housekeeping genes and pathways, such as transcription, translation, and proteostasis were significantly altered. In addition, over 150 genes and pathways that impact metabolic functions were altered, including imbalances in amino acid and protein degradation, dysregulated lipid metabolism, increased ketogenesis, and depletion of pathways involved in the metabolism of plant-based polysaccharides.

Regarding PD-relevant changes, we observed changes in pathways and genes that result in reduced levels of neuroactive molecules glutamate and serotonin, and mapped them to depletion of *Faecalibacterium prausnitzii*, *Blautia wexlerae*, and unclassified species. Evidence for increased toxicity included elevated synthesis of toxic trimethylamine (TMA) and increased degradation of beneficial dietary antioxidants, nicotinamide and amyloidogenic inhibitors, originating primarily from increased abundances of *Bifidobacterium* spp, *Acidaminococcus intestini*, and *Klebsiella pneumoniae*. We confirmed, via gut metagenomics, the finding in serum that secondary toxic bile acids are elevated in persons with PD ^31^, mapping it to *Clostridium scindens* and unclassified species. We confirmed and expanded on evidence for increased inflammation and pathogenicity. In addition to previously reported genes and pathways that regulate biosynthesis and immunomodulatory modification of lipopolysaccharides (LPS), cell wall, and lipoteichoic acid (LTA) cell wall ^22^, here we detected shifts in abundances of genes that are linked to virulence including regulators of quorum sensing, biosynthesis of toxins produced by pathogens, multidrug efflux pump, drug and antibiotic resistance genes, and reduced flagellin biosynthesis suggesting decreased capacity to stimulate beneficial mucosal immunity and promote enteric nervous and endocrine functions ^32^. Immunogenic features that are elevated in PD can be traced to opportunistic pathogens, including *Eisenbergiella tayi* and *Streptococcus mutans*. Protective features that are depleted, such as flagellin ^33^, map to a variety of species, including *Roseburia* spp.

To our knowledge, this is the first finding of disruptions in the CRISPR system and genes involved in DNA uptake and horizontal gene transfer in the PD microbiome. Changes in these genes, even in a few common taxa, can render the rest of the microbiome susceptible to bacteriophages, affect gene exchange, and alter the structure of the microbiome ^34^. Genes for Type-III CRISPR systems, which target and degrade RNA viruses, were reduced, whereas genes in Type-II CRISPR systems, Cas9, which target DNA, were elevated. Type-III genes map to *Roseburia intestinalis* and *Faecalibacterium prausnitzii*. Cas9-related genes map to *Bifidobacterium* and *Acidaminococcus* species. We also detected significant increases in competence protein genes, which facilitate the uptake of DNA, protect it from degradation, and help its integration into cellular DNA. Competence protein genes were encoded primarily by *Lactobacillus*, *Streptococcus,* and *Acidaminococcus* species. The IS6 family of transposase, which facilitates horizontal gene transfer, such as insertion of antibiotic resistance genes, was also enriched. IS6 mapped to *Klebsiella pneumoniae* and *Corynebacterium* spp.

Adding species of origin to the repertoire of genes and pathways for functional inference improved the information content of data. Noting species of origin ties in species’ differential abundances with functional inference. It revealed that depletion of *Roseburia intestinalis* and *Faecalibacterium prausnitzii*, whose detrimental effect has generally been attributed to reduced SCFA, also impairs the CRISPR system, and contributes to loss of glutamate and serotonin. We observed elevated tyrosine decarboxylase, which we might have inferred as clinically significant had we not mapped the species. Tyrosine decarboxylase made by the bacteria metabolizes L-dopa (a PD medication) into dopamine prematurely in the gut before it reaches the brain, thus attenuating its efficacy. However, the species of origin of this tyrosine decarboxylase gene was *Methanobrevibacter smithii* (an archaeon, not a bacterium), which uses the enzyme to turn tyrosine into tyramine. It is generally believed that SCFAs are depleted in the gut, but evidence is puzzling. Measured directly, SCFAs are reduced in stool but are elevated in the serum of people with PD ^35,36^. In the latest meta-analysis ^15^, fiber-degrading species that make SCFA were reduced, but pathways in the production of SCFAs were surprisingly elevated. We had a similar result, a drastic reduction in fiber-degrading species such as *Blautia* and *Roseburia* spp, but SCFA metabolism pathways had mixed results depending on the species of origin. Genes and pathways involved in fiber fermentation and SCFA metabolism/transport that map to *Blautia* and *Roseburia* spp were reduced, but SCFA-producing pathways that are encoded by *Bifidobacterium longum*, *Collinsella aerofaciens,* and unclassified species were elevated. We speculate that loss of fiber metabolizing taxa may have allowed *Bifidobacteria* spp to increase in abundance and fill a niche in SCFA production, although their repertoire, transport, and distribution (e.g., feces vs. serum) may be different.

Data extracted here are limited by what is known through culturable species, sequenced genomes, and characterized proteins. Much of the microbiome is still uncharacterized. In addition, metagenomics is entirely based on DNA; gene expression and metabolite availability are unknown. These limitations affect what we can or cannot detect, as well as our overall inferences about function. They do not, however, affect the differences we see in PD vs. NHC in what is detected. Thus, our results should be robust, although far from complete. Microbiome is still a very young field, with much to be learned and mined.

### Human genome and gut microbiome

PD has a strong genetic component. Genetic variants that increase the risk of PD are not fully penetrant, hence the notion that common forms of PD are caused by the interaction of genetic predisposition and environmental triggers, although the triggers and nature of interactions remain elusive. We questioned whether there is an interaction between the human genome and the gut microbiome in PD. Testing genome-microbiome interaction with an agnostic omics-wide approach is infeasible due to unattainable sample size and power requirements. We therefore opted for a hypothesis-driven study design that maximized power. We chose *SNCA* from the human genome and the clusters of fiber-degraders (20 correlated species) and opportunistic pathogens (16 correlated species) from the gut microbiome (**Extended Data 7-*SNCA***) to test interaction, pooled the three datasets to maximize sample size, and conducted the analyses using microbe set enrichment analysis (MSEA) ^37,38^. The strongest genetic risk factors for PD, with the highest peak in GWAS, are non-coding single-nucleotide polymorphisms (SNPs) that map to the *SNCA* gene region. Twenty unique SNPs at the *SNCA* locus have been associated with PD in GWAS ^39^. Based on linkage disequilibrium and correlation, we grouped the 20 SNPs into nine haploblocks; that is, they represent no more than 9 independent PD-associated signals (**Figure 3. *SNCA*-Microbe, Extended Data 7-*SNCA***). We show here that SNPs in four of the nine haploblocks (haploblocks 3, 4, 5, and 7) influence the dysbiosis in the PD gut.

**Figure 3.**
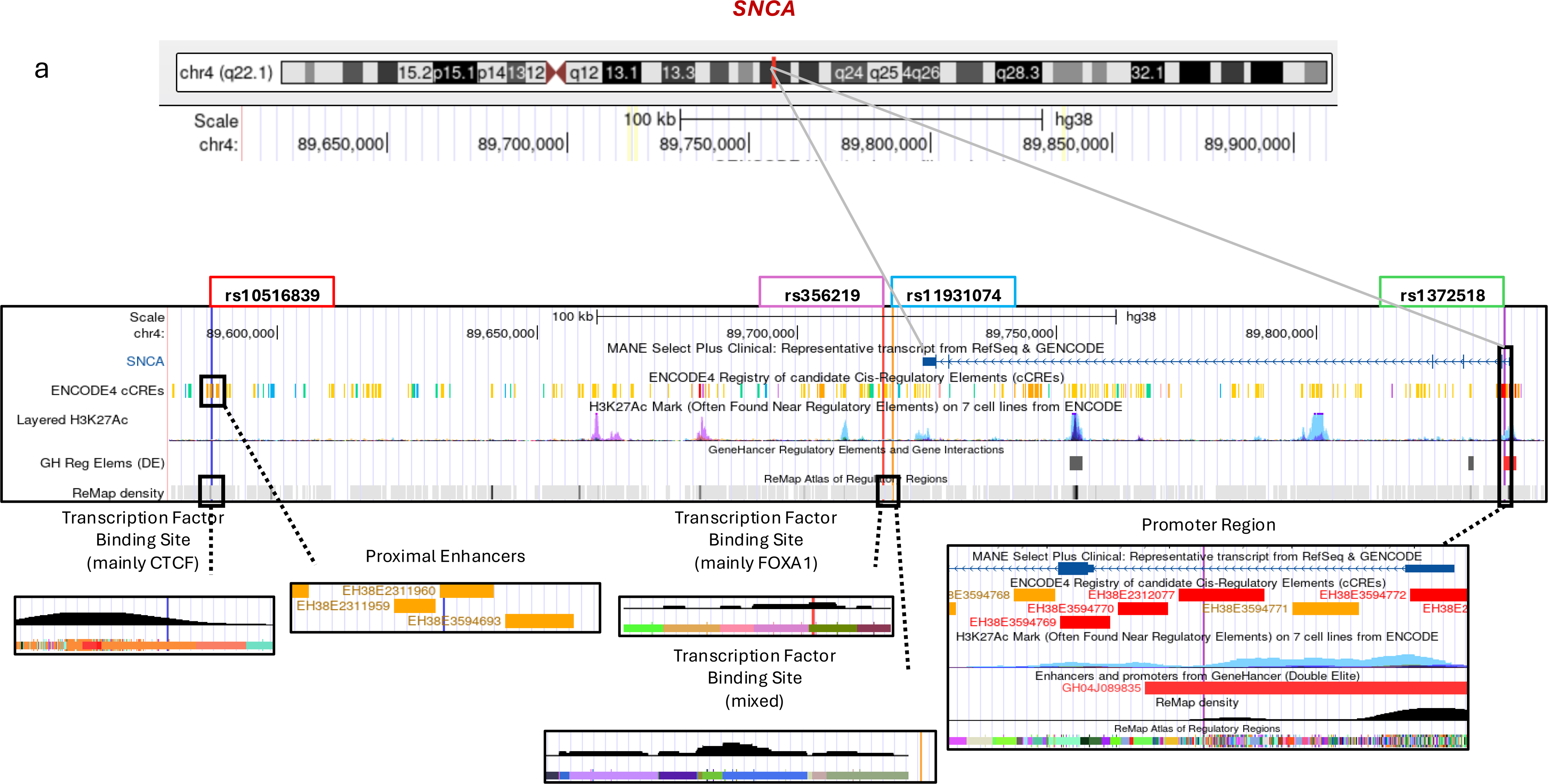
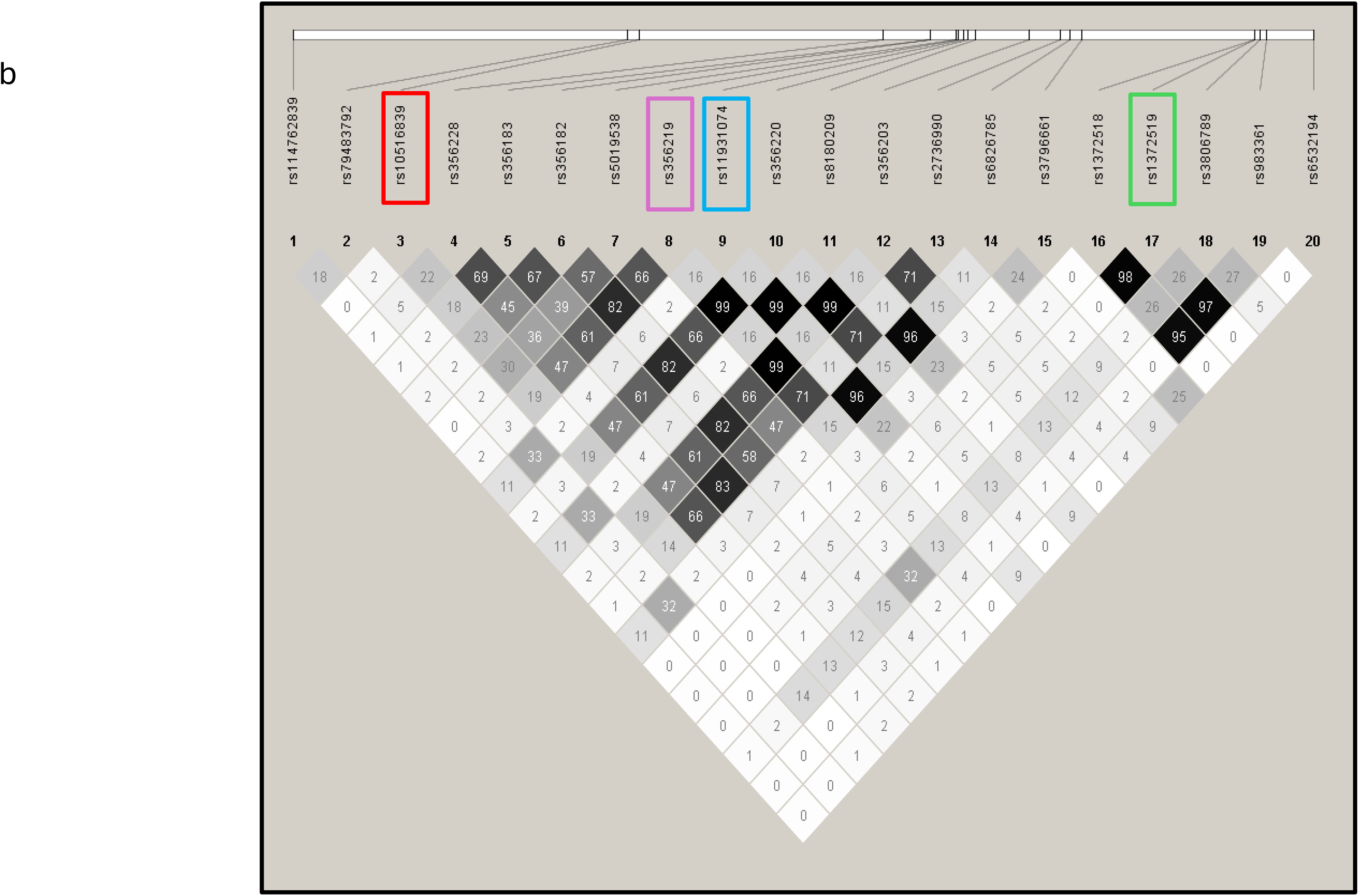
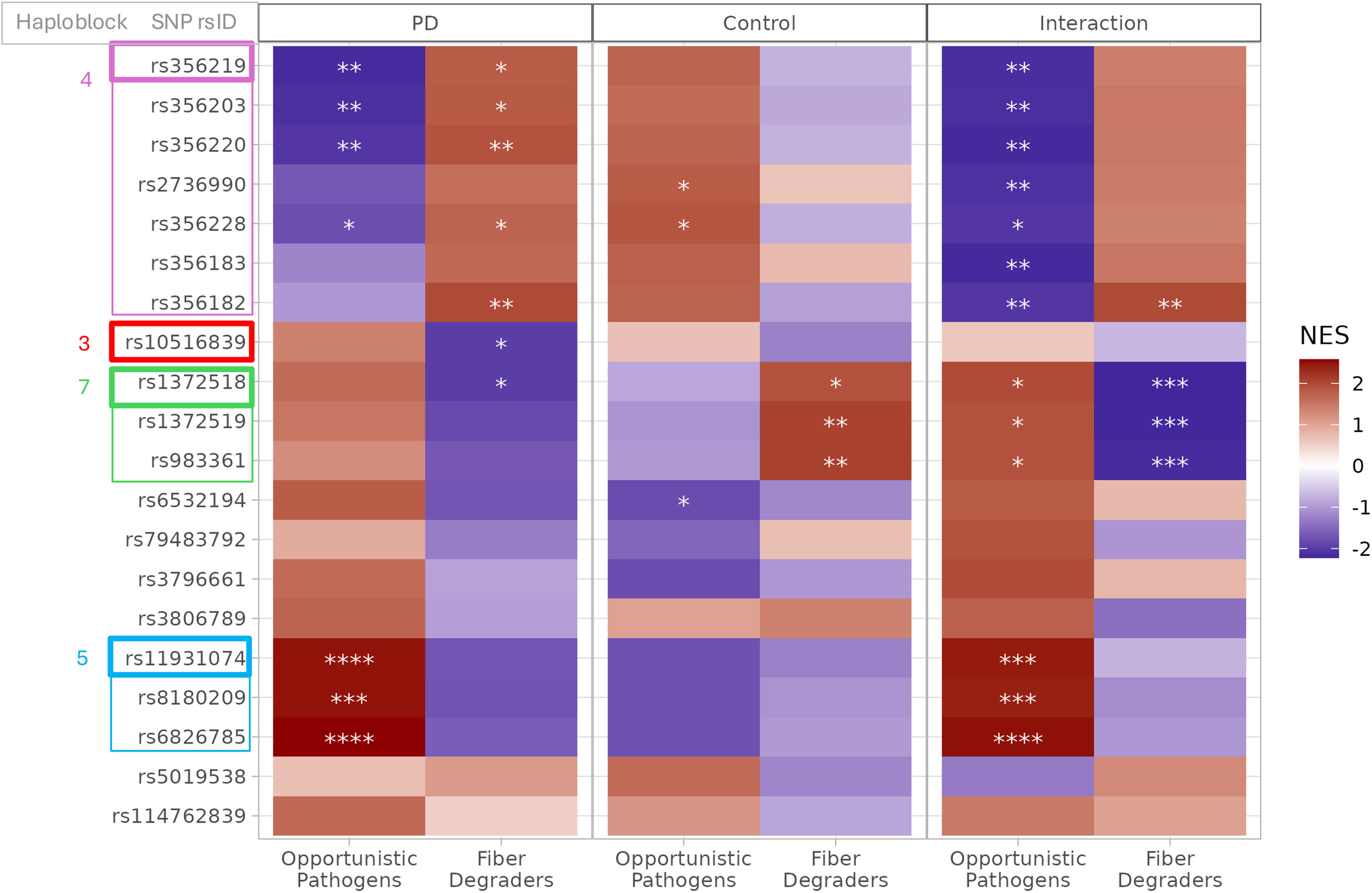
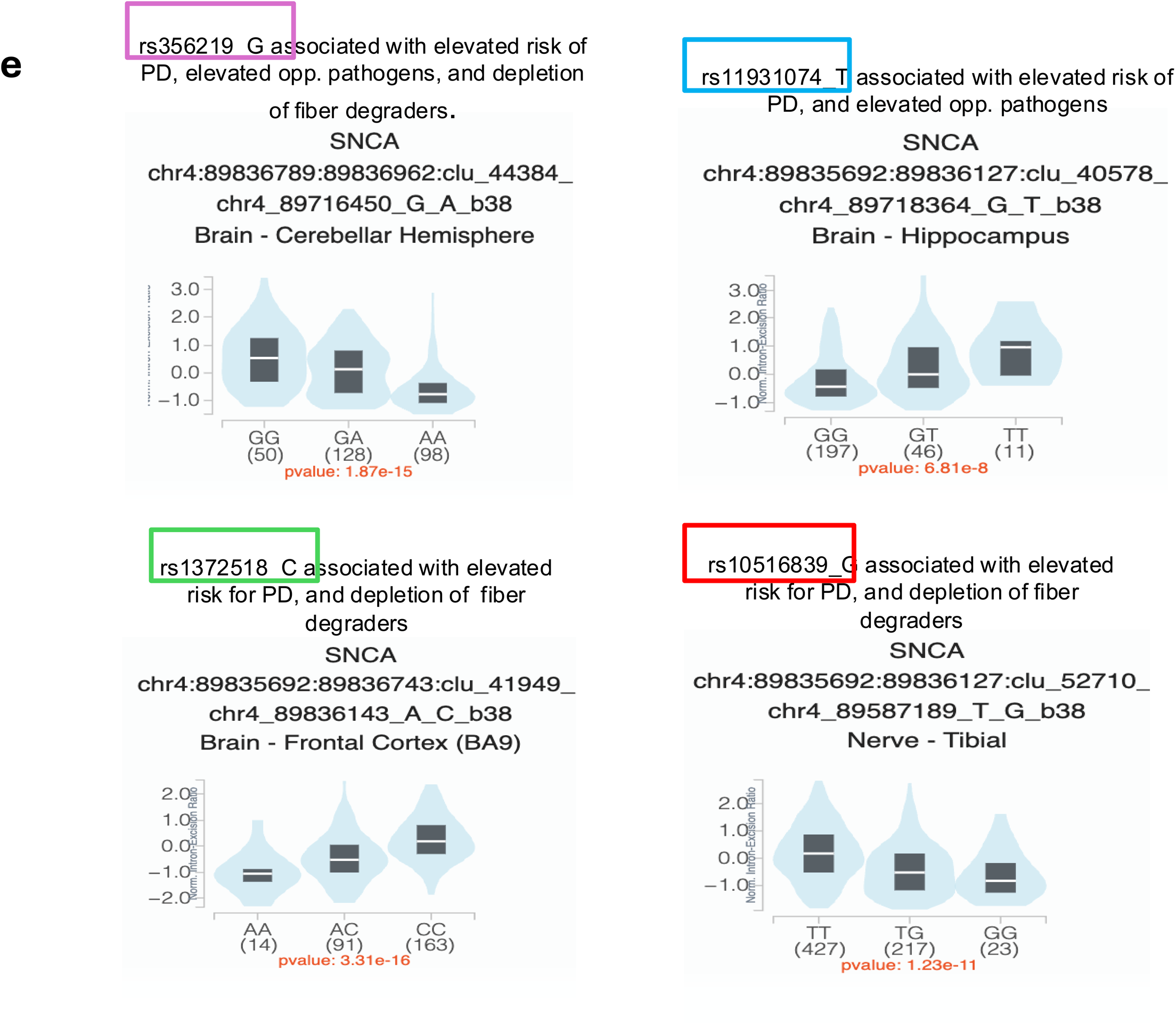
*SNCA*-Microbiome. PD-associated *SNCA* variants interact with the dysbiotic features in the PD gut microbiome. *SNCA* encodes alpha-synuclein, the pathological hallmark of PD. GWAS have identified 20 variants (SNPs) that map to *SNCA* region and are associated with increased risk of developing PD. PD-associated SNPs at the *SNCA* region were tested for interaction with dysbiotic features of PD (Table 4. SNCA-microbe). SNPs from four haploblocks were associated with differential microbial abundance. Here, we present data on four SNPs, representing the four haploblocks that interacted with microbial features. The four SNPs are in very tight linkage with other SNPs in their haploblock and produce nearly identical results. There is no correlation across the four blocks (r2 = 0 to 0.19); they are independent. The SNPs are indicated in pink (haploblock 4), red (haploblock 3), blue (haploblock 5), green (haploblock 7) a. Top: Position of the *SNCA* gene on chromosome 4. Middle: SNCA region zoomed out to show the position of PD-associated SNPs that interact with dysbiotic features of PD. One is in the promoter 5’ of SNCA, the other 3 are at the 3’ of the gene. Bottom: Further magnification showing SNPs are in regulatory enhancer regions. b. Haploview of all 20 PD-associated SNPs. Cells show the pair-wise correlation between the SNPs, which was the basis for assorting the 20 SNPs into 9 haploblocks. c. Heatmap depicting results of microbiome set enrichment analysis (MSEA). *SNCA* SNPs were tested for association with relative abundances of fiber degraders and opportunistic pathogens (**Extended Data 7-*SNCA***). In the heatmap, red indicates increased abundance, and blue indicates decreased abundance of microbe as a function of the number minor alleles. Star indicates significance at FDRx9, where FDR is multiple-testing correction performed by MSEA, multiplied by 9 as further Bonferroni correction for 9 independent haploblocks tested. FDRx9 * < 0.05, **<0.01, *** <1e-3, **** <1e-4. d. Data and figures were extracted from the Genotype-Tissue Expression (GTEx) database. SNPs from each haploblock were queried in the GTEx database and found to be sQTL, affecting the splicing of the SNCA transcript. Each SNP affects a different splice site. Splicing of the SN CA transcript results in four major alpha-synuclein isoforms with varying affinity for aggregation into pathologic form.

**Table 4.**
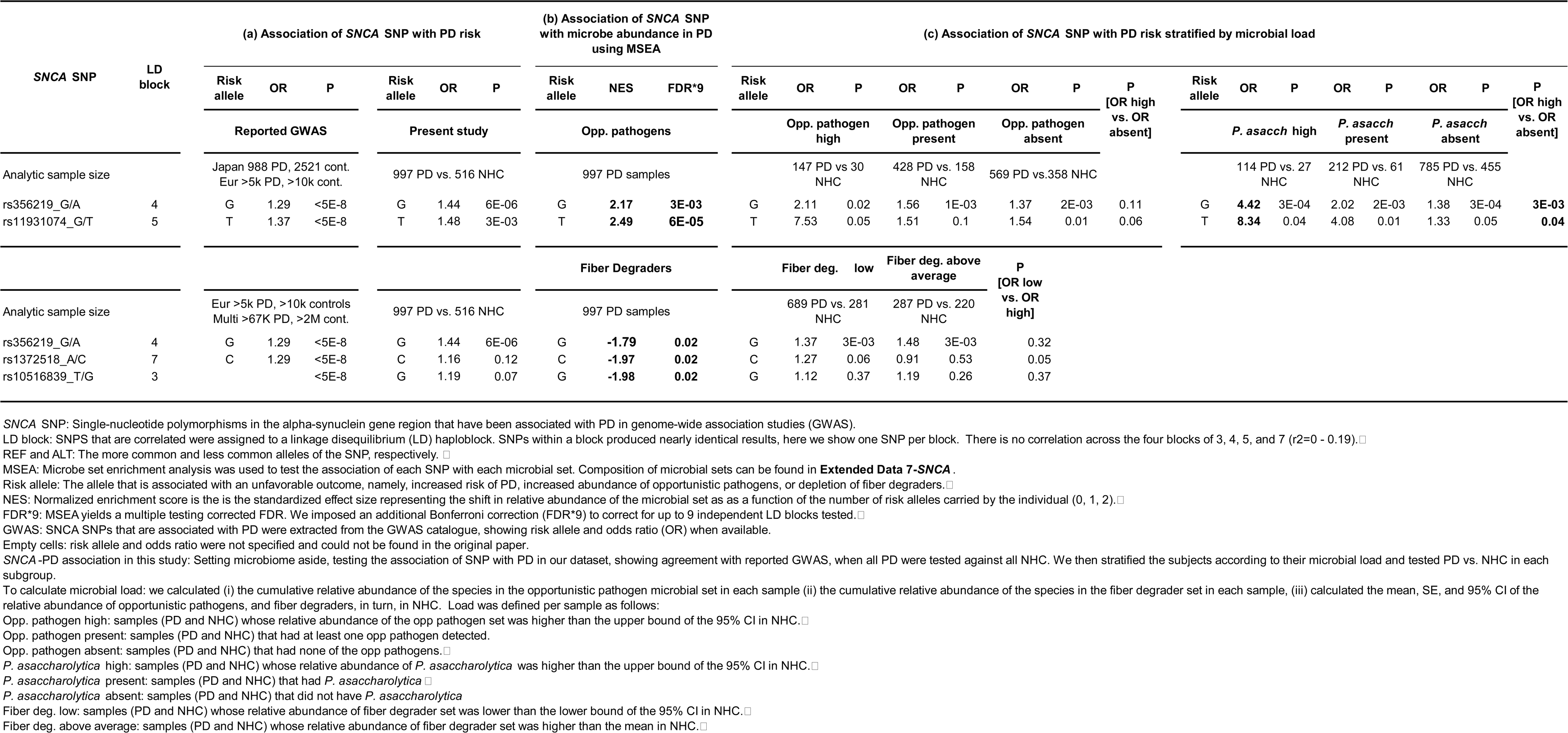
Genetic variants at the SNCA locus and the dysbiotic features of gut microbiome in PD. Investigating the effect of SNCA genotype on dysbiotic features of microbiome, and the effect of gut microbes on SNCA -associated genetic risk for PD. (a) PD-associated variants (SNPs) in the SNCA region were extracted from the GWAS Catalog. Odds ratio (OR) and risk allele were obtained from the GWAS Catalog and original papers when available, and were also calculated in the present dataset. (b) SNPs were tested for association with the relative abundance of opportunistic pathogens and fiber degraders in PD samples using the microbe set enrichment analysis method (MSEA). Full results are in Extended Data 7-SNCA. Shown are SNP-microbe associations that were significant. Note the directionality is consistent, i.e., of the two alleles of each SNP, the allele that increases PD risk is the same allele that associates with increased abundance of opportunistic pathogens and decreased abundance of fiber degraders. (c) SNCA SNPs were tested for association with PD risk (as done in GWAS), but now in subgroups stratified by microbial load. High levels of opportunistic pathogens, and Porphyromonas asaccharolytica specifically, were associated with increased penetrance of SNCA SNPs, elevating SNP-PD association from OR∼1.4 to OR∼4.5 for SNPs in block 4, and from OR∼1.5 to OR∼8 for SNPs in block 5.

First, we tested the association of *SNCA* SNPs with microbe sets in metagenomes of persons with PD, i.e., does genotype at *SNCA* affect the relative abundance of dysbiotic features in PD? (**Table 4. *SNCA*-Microbe, Figure 3. *SNCA*-Microbe**). We found that the elevation in relative abundance of opportunistic pathogens was associated with *SNCA* SNPs in haploblocks 4 and 5, and the reduction in fiber degraders was associated with SNPs in haploblocks 3, 4, and 7. The gene-microbe associations detected in the PD metagenome were absent in NHC metagenomes, which, coupled with highly significant tests of interaction, suggest these *SNCA*-microbe associations are specific to the disease microbiome and are not a general phenomenon.

As is evident in the MSEA heatmap (**Figure 3. *SNCA*-Microbe**), SNPs that were associated with an increase in opportunistic pathogens were often associated with a decrease in fiber degraders, and vice versa. Note, however, that because there was no correlation between the relative abundances of opportunistic pathogens and fiber degraders (**Figure 1. Networks**), the association of *SNCA* with each, in opposite directions, suggests the interactions are independent.

Importantly, directionality aligned consistently, i.e., of the two alleles of each *SNCA* SNP, the allele that was associated with an increase in opportunistic pathogens or a decrease in fiber degraders was the same allele that was associated with increased risk of PD in published GWAS and in the present dataset (**Table 4. *SNCA*-Microbe)**.

SNPs identified here are not in the coding regions of *SNCA* (**Figure 3. *SNCA*-Microbe**). To explore if they regulate the expression of *SNCA*, we used the GTEx database (https://gtexportal.org), a repository of genome-wide genetic variants that are associated with gene expression (expression quantitative trait loci, eQTL) or RNA splicing (sQTL). rs10516839 (block 3) is an eQTL for *SNCA* in the brain (P=9E-5); the others are also noted as eQTLs but not in the brain or the GI tract. Notably, SNPs from all four haploblocks are sQTL for *SNCA* in the brain, with high significance (P=7E-8 to 3E-16) and large effect sizes (**Figure 3. *SNCA*-Microbe**). Each SNP affects a different splice site. Splice variants generate transcripts with varying lengths that are translated into four major alpha-synuclein isoforms with different affinities for aggregation ^40^. Thus, the SNPs identified here could potentially contribute to pathology via the regulation of gene expression and the promotion of alpha-synuclein aggregation.

Second, we explored the notion that microbes may influence the penetrance of *SNCA* SNPs in their association with PD. GWAS have identified hundreds of susceptibility genes for common disorders, but these genes typically exhibit very low penetrance (for example, *SNCA* variants have an odds ratio (OR) of less than 1.5). A significant and unresolved question is which factors interact with these genetic predispositions to trigger the development of disease. To test if the microbiome harbors such triggers, we divided cases and controls according to the load of dysbiotic features and estimated the OR of the association of each SNP with PD in each stratum. For opportunistic pathogens, the strata were microbe absent (not detected), present (detected irrespective of relative abundance), and high (relative abundance in the individual higher than the upper 95% confidence interval (CI) boundary in controls). For fiber degraders, strata were low (relative abundance in the individual lower than the lower 95% CI boundary in controls) and above average (higher than the mean in controls) (**Table 4. *SNCA*-Microbe, Extended Data 7-*SNCA***). Fiber degraders did not influence the association of *SNCA* with PD risk. In contrast, opportunistic pathogens increased the penetrance of the PD-associated *SNCA* SNPs in haploblocks 4, and 5. Having high levels of opportunist pathogens combined as a cluster increased the association of rs11931074 (haploblock 5) with PD from OR=1.5 to OR=7.5, and *Porphyromonas asaccharolytica* as a single species of opportunistic pathogen, elevated *SNCA*-associated PD risk to OR=8.3. These data suggest opportunistic pathogens in the gut are a trigger for genetic susceptibility at *SNCA*.

This finding is novel in having identified a factor that elevates genetic risk by several-fold. In addition, the interaction with *SNCA* on PD risk suggests opportunistic pathogens in the gut are involved in disease causation, which is seldom possible to establish in human studies. The caveat is that the analysis was conducted on the entire dataset (1,006 PD and 544 controls), and although this is considered a very large sample size in microbiome studies, genetics necessitates independent replication.

Studies of the human genome and microbiome are often founded on the premise that disease genetics shapes the microbial dysbiosis, which can sometimes lead to disease ^41,42^. Here, we show that the reverse can also be true, that microbes can increase the penetrance of genetic risk. The premise of the disease gene leading to dysbiosis can go in two directions: one, the *SNCA* variant elevates pathogens, and elevated pathogens increase the risk of PD, or two, the *SNCA* variant is a risk factor for PD, and having PD allows for pathogens to grow. The former is unlikely because if true, the *SNCA* variants would be associated with elevated pathogens in NHC as well, and they are not. The latter is also unlikely because if PD is causing the increase in pathogen abundance, then why is pathogen abundance associated with specific *SNCA* variants, and not all PD-associated variants, or even all PD irrespective of genetics? Our data are most consistent with a synergistic interaction of pathogens with specific *SNCA* variants, which together elevate the risk of PD higher than the *SNCA* variant alone. Literature in this area has noted *Porphyromonas gingivalis* in the brains of Alzheimer’s disease, and that *P. gingivalis* has a toxic effect on tau via virulent cystine proteases ^43^. We do not see involvement of *P. gingivalis* in PD, but rather *P. asaccharolytica*. Moreover, the microbe-associated SNPs in PD do not encode amino acids of alpha-synuclein; they are non-coding variants that regulate the expression and splicing of the *SNCA* transcript. Other lines of evidence, unrelated to PD, have shown that upon infection with a pathogen, *SNCA* expression is turned on as a necessary step to induce an immune response to fight infection ^44,45^. We suggest that this process could set the stage for future development of PD, and more so for individuals who carry *SNCA* SNPs that yield aggregation-prone isoforms of alpha-synuclein.

### Microbial features as biomarkers for subtyping

Microbiome-based clinical trials for PD therapies are on the rise. Results of early clinical trials with pre/probiotic supplementation ^27–29^ and fecal microbiome transplant (FMT) ^46,47^ are mixed but promising. Next-generation targeted therapeutics are forthcoming. New therapies for PD are desperately needed, and clinical trials offer hope to patients and their families. Most clinical trials recruit only patients in early stages of disease, leaving families with late-stage PD feeling abandoned. Microbiome-based therapies have the potential to help patients at any stage of disease by bringing symptom relief and slowing disease progression, even if it is too late to address the root cause of the disease.

PD is heterogeneous. All disease-modifying drugs for PD have failed in clinical trials, in part because, without biomarkers, trials cannot select the appropriate patient population for their target. Can we do better with microbiome-based clinical trials using dysbiotic features as biomarkers? Here, we present a first view of the metagenomic heterogeneity in PD gut, present a conceptual framework to define subtypes, and argue for metagenomic prescreening to enhance the power of clinical trials for microbiome-based therapeutics. The concept of prescreening to enrich trial cohorts is not novel: high blood pressure drugs are tested in people with high blood pressure, PD drugs targeting the *LRRK2* G2019S mutation would not include patients who do not have this mutation. The novelty of this study is in demonstrating the inter-individual heterogeneity in PD gut microbiome, the ability to identify the subtypes, and the feasibility of using dysbiotic features as biomarkers for prescreening and enrichment of clinical cohorts (**Figure 4. Subtypes).**

**Figure 4.**
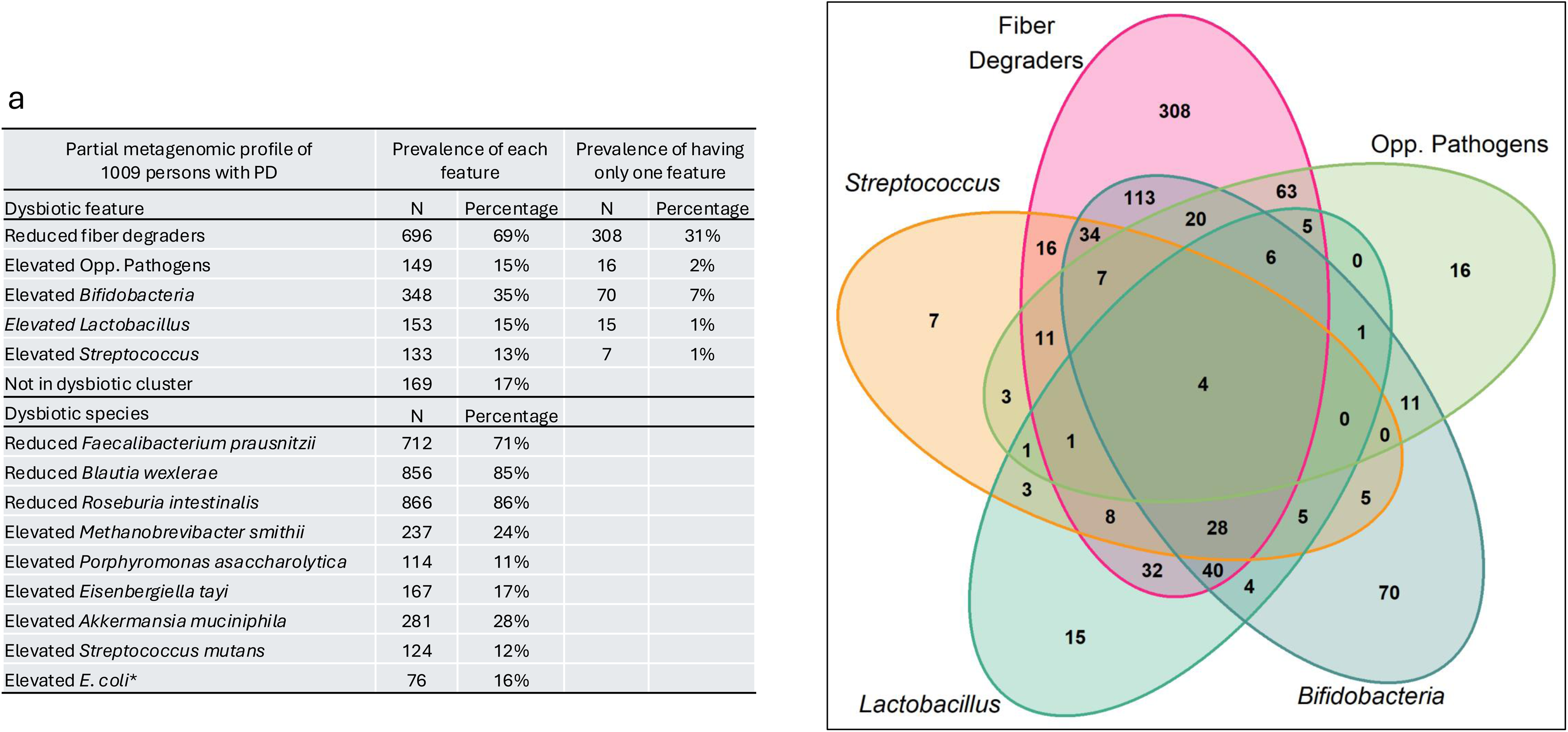
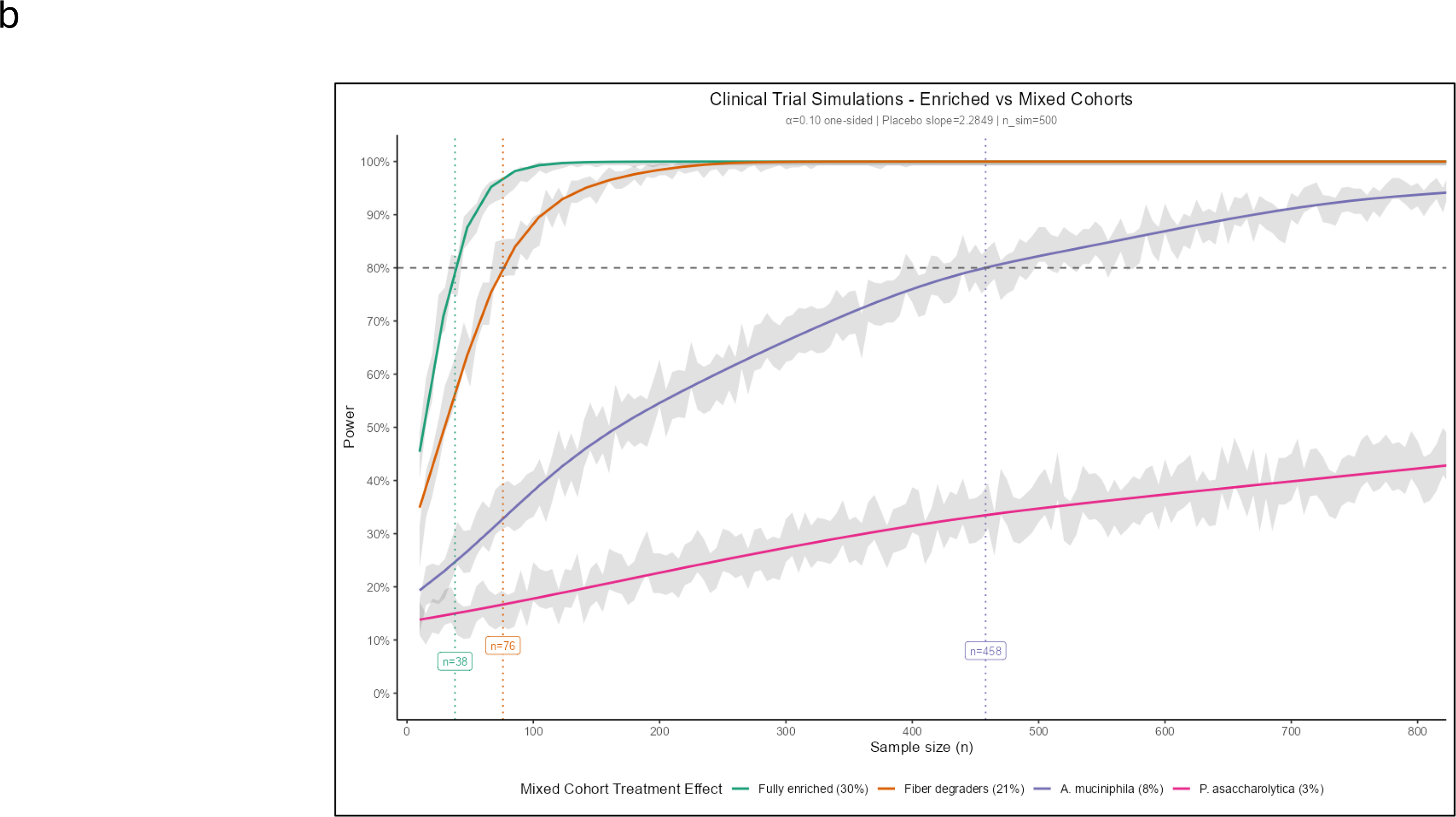
PD subtypes defined according to dysbiotic features in the gut. Data presented show that not all persons with PD have a dysbiotic microbiome, and not all dysbiotic guts have the same features. Metagenomic profiles of 1006 persons with PD were used to demonstrate the inter-individual heterogeneity of dysbiotic features, once based on five PD-associated polymicrobial clusters, and again according to seven PD-associated species. These are for demonstration. Classification can be done for any target of interest using individual-level relative abundances presented in Extended Data 8-Subtyping. (a) Number and percentage of patients with each dysbiotic polymicrobial feature. The overlaps indicate the number of patients with more than one dysbiotic feature. E.g., 696 (69%) of patients had low fiber degraders; among them, 308 (31% of all patients) had only low fiber-degraders with no other dysbiotic feature detected, and 338 (38% of patients) had other dysbiotic features in addition to depletion of fiber-degraders. Subtyping at the polymicrobial level can aid subject selection for current treatments such as FMT and pre/probiotics. Next generation targeted therapeutics will require higher resolution subtyping by species or microbial genes. Here we show 71% of patients have low *Faecalibacterium prausnitzii*. *Akkermansia muciniphila* is elevated in 29% of patients. Pathogenic species *Eisenbergiella tayi*, *Sterptococcus mutans*, *Porphyramonas asaccharolytica*, and *E. coli* are each elevated in less than 20% of patients. (\**E. coli* prevalence is based on DOD dataset (487 PD patients)). (b) Power and sample size simulations demonstrate that prescreening and enriching the cohort for the dysbiotic feature increase the efficacy of clinical trials. Data were generated using parameters specified in https://clinicaltrials.gov/study/NCT04658186: treatment effect=30% (measured as placebo effect in untreated arm being 2.285 increase in UPDRS I-III score/year, and 30% improvement being (2.285*1-0.3)=1.59 increase in UPDRS I-III score/year in treatment arm), power=0.8, alpha=0.1. In this scenario, if patients are prescreened and the cohort is enriched for the dysbiotic feature, 38 patients each in treatment and placebo arms will have 80% power to detect the 30% treatment effect (green line). Orange, blue, and pink lines represent the dramatic loss of power if cohort is not enriched and the dysbiotic feature is present only in a subset of patients. Orange line: when the feature is present in 70% of patients (e.g., fiber degraders) the treatment effect dilutes to ∼20% for the cohort overall, doubling the required sample size to 76 patients in each arm. Blue line: when the feature is present in 30% of patients (e.g., *Akkermansia muciniphila*), the average treatment effect for the cohort drops to about 8%, and the sample size to detect it escalates to over 400 patients in each arm. Pink line: when the feature is present in only 10% of the cohort (e.g., *Porphyromonas assachrolytica*), the average treatment effect becomes so low, ∼3%, that a sample size of 800 patients will yield only 30% power. This graph shows clinical trials for uncommon features are not futile if patients are prescreened and the cohort is enriched in the feature of interest, effectively raising the pink, blue, and red lines up to the green line.

Metagenomics studies, including the sections above, assess the overall differences between a PD population and a non-PD population. Here, we turn the approach around, from the population level to the individual level, to ask what the microbiome of an individual patient looks like. Do all people with PD have dysbiotic microbiomes? Do they all have the same features, or is there heterogeneity in dysbiotic features? Using individual-level taxonomic relative abundance data (**Extended Data 8-Subtyping**), we can classify patients by their dysbiotic features, thereby defining microbiome-based subtypes of PD. To do this, we used the distribution of relative abundances of species in controls as the reference framework and classified a PD metagenome as dysbiotic if it had very high or very low levels of PD-associated features (outside 95% CI boundaries of controls, see **Methods**). Here, we demonstrate the concept with polymicrobial features, which would be more applicable to non-specific treatments such as diet, pre/probiotics supplementation, and FMT, and with specific species or genes, which would be necessary to guide next-generation targeted therapeutics. In practice, the feature would be the target of the drug being tested in a clinical trial. It will require a reference dataset (controls) to gauge the frequency distribution of the feature of interest, which will then inform pre-screening of potential participants to select the optimal patient population for the clinical trial.

First, we used the DOD dataset to define six PD-associated polymicrobial features, applied it to all three datasets to estimate their prevalences, and found reasonable consistency across datasets in the prevalence of the dysbiotic features (**Extended Data 8-Subtypes**). Next, we pooled the three datasets. We used results of MWAS meta-analysis to select a few PD-associated features: fiber degraders, opportunistic pathogens, *Bifidobacteria* spp, *Lactobacillus* spp, and *Streptococcus* spp. We calculated the frequency distribution of each feature in NHC with 95% CI boundaries. We then turned to PD samples. For features that are enriched in PD vs controls, a metagenome was tagged as dysbiotic if the relative abundance of the feature was greater than the upper bound of the 95% CI in NHC. Similarly, for fiber degraders that are depleted in PD, a metagenome was tagged as dysbiotic if the relative abundance of the feature was lower than the lower bound of the 95% CI in NHC. Having identified each feature of interest in each patient, we tabulated the counts and estimated the prevalence of each feature and its co-occurrence with other features (**Figure 4. Subtypes**). Among polymicrobial clusters, the most common dysbiotic feature was depletion of fiber degraders, seen in 69% of patients.

Treatments with diet and prebiotics or probiotic supplementation to restore this deficiency may be beneficial to many patients. They may even help balance the abundance of species, such as *Akkermansia muciniphila,* that are negatively correlated with fiber degraders. However, half of the patients with depleted fiber degraders have other dysbiotic features that are not correlated with fiber degraders, such as opportunistic pathogens and *Streptococcus* spp. When dietary and pro/prebiotic supplementation is the treatment (most current trials listed in clincaltrials.gov), prescreening to exclude patients with co-occurring dysbiotic features could improve the odds of achieving meaningful clinical improvement. Elevated *Bifidobacteria* spp was the second most common polymicrobial feature, affecting 35% of patients. Other features were less common: 15% of patients had high opportunistic pathogens, 15% had high *Lactobacillus* spp, and 13% had high *Streptococcus* spp. Notably, 17% had none of these features, a group that should be identified and excluded from any microbiome-based trial. To demonstrate the dramatic gain in power that could be achieved by prescreening and enriching the cohort with patients who have the targeted dysbiosis, we present simulated power and sample size calculations in **Figure 4. Subtypes**.

Next-generation targeted therapeutics will require higher resolution than polymicrobial groups. Here, we show the prevalence of species that are commonly noted in PD studies. A similar framework can be obtained for microbial genes. Over 70% of patients have very low levels of fiber-degrading species *Blautia wexlerae*, *Roseburia intestinalis*, or *Faecalibacterium prausnitzii*. *Akkermansia muciniphila* is elevated in 28% of patients, *Methanobrevibacter smithii* in 24%, *Eisenbergiella tayi* in 17%, *E. coli* in 17%, *Streptococcus mutans* in 12%, and *Porphyromonas asaccharolytica* in 11%. Prescreening patients will be critical to the success of targeted therapies if only 10%-30% of patients can respond (**Figure 4. Subtypes)**.

Many of the definitions and threshold sets in this proof-of-concept exercise are arbitrary and can be changed to better suit a given study. Although we found similar profiles in datasets from northern and southern US, the composition and prevalence of dysbiotic features could vary globally, requiring a region-specific reference framework. Owing to a strong push for open science, most studies worldwide now share their raw individual-level metagenomics data publicly, which is sufficient to recreate the reference framework. This work is intended as a conceptual presentation, a starting point to explore the notion of using microbiome profiles as biomarkers, as a diagnostic companion for patient selection for microbiome-based clinical trials, and, if successful, for individualized treatment.

In conclusion, this large-scale, three-pronged study highlights the potential of the microbiome to uncover microbial contributors to disease, links genetic predisposition to microbes that elevate disease risk, and demonstrates the utility of dysbiotic features as biomarkers for microbiome-based clinical trials and personalized therapeutic strategies.

## Methods

We complied with ethical regulations approved by the Institutional Review Board (IRB) for Protection of Human Subjects at the University of Alabama at Birmingham (UAB). All subjects gave signed informed consent. No compensation was made for participating in the study. For a schematic workflow, see **Extended Data 1-STORMS**.

### General rules

Software and reference databases used in the study are named in the text; the versions and the URLs are provided in **Table 5-Key Resources**. Statistical significance was set at FDR < 0.05 for when multiple testing occurred, FDR < 0.05\**N* if analysis was repeated in *N* number of independent tests, and P<0.05 when uncorrected permutations were used.

**Table 5.**
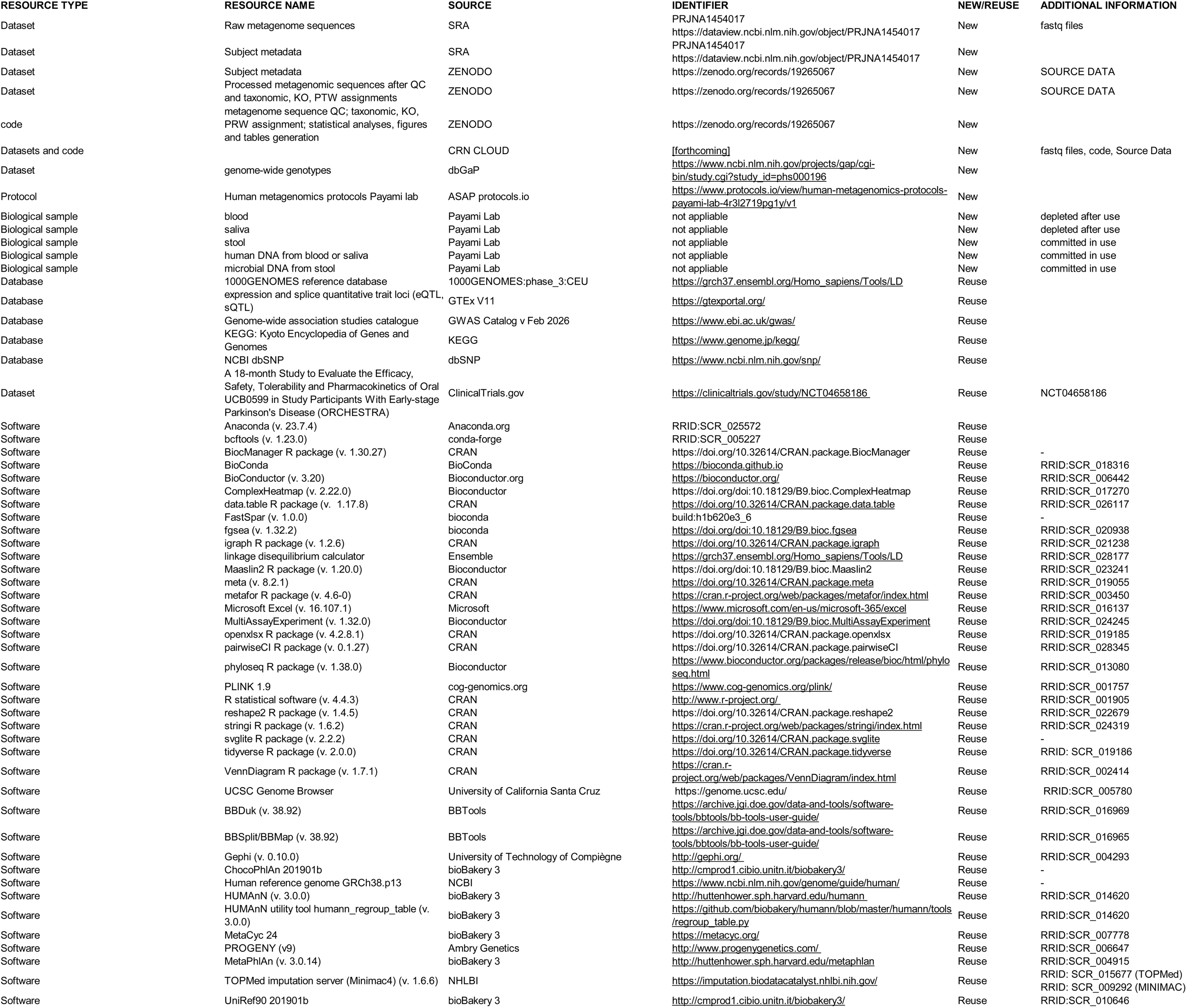
Key Resources.

Statistical testing was done on each dataset separately, then meta-analyzed using a random effects model. If heterogeneity across datasets was significant (I^2^ ≥ 50%, P_Het_ < 0.1), the result of the meta-analysis was deemed unreliable as a true representative of the underlying datasets. A feature was called significant in meta-analysis if FDR<0.05 and P_Het_>0.1. Every analysis was performed by two analysts and double-checked. The data and code are available on NCBI SRA and Zenodo (**Data and code availability**).

### 1. Standardization

Microbiome studies of PD are highly vulnerable to methodological variation ^15^. It is therefore a paramount advantage to keep variation to a minimum. This study, having been conducted by the same investigators since the start, was standardized to every degree possible. The study started in 1991 with enrolling subjects for a genetic study of PD who were later enrolled in the microbiome arm. The microbiome arm was added in 2014, and three microbiome datasets were collected over 6 years (2014-2020). Unless noted, methods apply uniformly to all datasets throughout the study, from subject selection to the statistical pipeline. Protocols were kept untouched except when updated to incorporate new knowledge or advances in the field.

Changes were made only as new pertinent data were released (e.g., added Bristol Stool Chart and medication to metadata for DOD), or when a better method was developed, and only at the start of each dataset. We kept detailed records down to the slightest deviations to track its potential consequences.

There was one unavoidable change during the study that had considerable consequences: the DNA collection kit. NGRC and UAB recruitment started in 2014 and 2015, respectively, for which we used sterile swabs for stool collection, the best choice at the time, barring immediate freezing, which is not feasible for large-scale studies. When we launched, we switched to the OMNIgene*GUT collection kit, which had proved in comparative studies to be superior to swabs and other preservatives. Stool samples were collected at home and mailed to the lab, and immediately frozen. During shipment at ambient temperature, *E. coli* and *Klebsiella* spp had grown in PD and control samples alike. We retained *E. coli* and *Klebsiella* spp in the NGRC and UAB datasets, as their exclusion would have further altered the overall composition. Instead, we considered results on *E. coli* and *Klebsiella* spp from NGRC and UAB unreliable. Any results presented on *E. coli* or *Klebsiella* spp are based on the DOD dataset.

### 2. Data integrity and Reproducibility

To ensure data accuracy and reproducibility, quality control steps were incorporated as follows. Data collection questionnaires that were to be self-administered by the subjects had redundant data to cross-check for mix-ups and inaccuracies. Upon receipt of each packet, the two questionnaires, stool sample, and saliva sample in the packet were cross-checked for potential labeling error in the lab, sample mix-up at home, which occurs when a patient and a control are in the same household, for completeness, comprehensibility, and lucidity of information provided. Discrepancies were either resolved by checking the medical records and calling the subject, or the sample or metadata in question was excluded. Data from questionnaires were entered in the database twice, by two individuals, one in a customized PROGENY database, and one in Excel spreadsheets, downloaded, cross-checked for 100% of entries, errors identified and corrected, and repeated cross-checks until clean. All bioinformatic and statistical analyses were conducted by two analysts using the same or different packages but writing their own codes; the results were compared, inconsistencies identified and corrected. The senior author oversaw every step, from patient selection to data analysis outputs, reviewed the input and outputs, caught and helped correct inconsistencies that could escape analysts’ and technicians’ notice, and ensured consistency and adherence to standardization. To ensure others can reproduce these results, we made the raw and processed data publicly available on the SRA and Zenodo repositories, provided the step-by-step workflow and inclusions and exclusions for enrollment and metadata, metagenome, and genotypes in STORMS charts, and the code created and used for bioinformatics and statistical analyses on Zenodo.

### 3. Generating the data

#### 3.1. Recruitment of Study participants (Extended Data 1-STORMS)

In 2014, when we started a pilot study on microbiome, we had a large cohort of PD and NHC collected since 1992 and had been used in GWAS ^48^. We recontacted the surviving participants and enrolled 210 PD and 134 NHC subjects into our pilot NGRC microbiome study. NGRC subjects were from the states of Washington (45%), New York (42%), and Georgia (13%). The second dataset was collected during the 2015-2017 period from the University of Alabama at Birmingham (UAB) and included 309 PD and 178 NHC subjects from Alabama and surrounding states in the Deep South. The third dataset was also collected at UAB with funding from the Department of Defense (DOD) and included 487 PD and 232 NHC enrolled from 2018 to 2020. Patients were enrolled from neurology clinics in four states (Washington, New York, Georgia, and Alabama). Patient recruitment was sequential irrespective of disease stage, family history, or any other risk factor. The only inclusion criterion was diagnosis of PD by a movement disorder specialist. Controls were spouses of PD patients, or community volunteers of similar age (preferably over 50 years of age), who did not have neurologic disease by self-report (no PD, Alzheimer’s disease, dementia, multiple sclerosis, amyotrophic lateral sclerosis, ataxia, dystonia, autism, epilepsy, stroke, bipolar disorder, or schizophrenia). Controls were from the same geographic areas as patients. Each potential participant was met by a study staff member in private, the study and their involvement in it were explained, they were given time to read the Informed Consent Form and ask questions, and if they agreed to participate, sign the consent. After signing the informed consent, subjects were given a packet containing two questionnaires (metadata), a stool collection kit, a saliva collection kit if they had not donated blood, detailed instructions, and a self-addressed pre-stamped envelope for return. Samples donated after the start of the COVID-19 pandemic in the US were not used in the study.

#### 3.2. Metadata

were collected using two self-administered questionnaires and reviews of medical records. The two questionnaires were the Environmental and Family History Questionnaire (EFQ) for PD-related data, and the Gut Microbiome Questionnaire (GMQ) for gut health, which was completed immediately after stool collection. EFQ and GMQ were completed by the subject without investigator interference. Electronic medical records were used to extract or verify clinical data by a movement disorder specialist neurologist.

#### 3.3. Stool samples

Each subject provided one stool sample at one time point. Each data point is a unique individual. Each sample was measured and analyzed once. Stool samples were collected at home and mailed back to the lab by the US Postal Service. For the NGRC and UAB cohorts, which began collection in 2014 and 2015, respectively, we used Fisher Scientific (Hampton, NH, USA) DNA/RNA-free BD BBL Sterile/Media-free Swabs. For the DOD cohort, we used DNA Genotek (Ottawa, Ontario, Canada) OMNIgene*GUT collection kit, which has a preservative (see **Standardization** section). Upon receipt in the lab, samples were immediately examined and cross-checked for QC and stored in -20°C freezer.

#### 3.4. DNA extraction from stool and next-generation metagenome sequencing

Case and control samples were randomly assigned and interspersed on plates to minimize batch effect. Positive and negative controls and technical replicates were also included on plates for extraction and sequencing. DNA was extracted using the same chemistry with the PowerSoil and PowerMag kits from MO BIO Laboratories (Carlsbad, CA, USA), which was acquired by Qiagen in 2016 (Germantown, MD, USA). Library preparation was done using the Illumina Nextera system and sequenced on Illumina NovaSeq 6000 instrument, aiming for 40-50 million reads per sample, following a paired-end 150 base pair (bp) sequencing protocol. DOD samples were extracted and sequenced by CosmosID (Germantown, MD, USA), achieving a mean of 50 million raw sequences per sample, reduced to 35 million after QC. For NGRC and UAB, DNA was extracted by the Rob Knight Lab at the University of California, San Diego ^49^, and later sequenced at Zymo, Inc., achieving a mean of 62 million raw sequences per sample, reduced to 40.4 million for NGRC and 36 million for UAB after QC.

#### 3.5. Genotyping the human genome

Subjects donated a blood sample or saliva for genetic studies. DNA was extracted from whole blood or saliva, and genome-wide genotyping was conducted on the Illumina genotyping array platform ^48^. Genome-wide genotyping and subsequent QC were conducted at Johns Hopkins Genetic Research Core Facility, using standardized protocols for GWAS, and generating an average of 1.2 million hard call genotypes per individual. Imputation was conducted in 2022 using the TOPMed imputation server ^50^ (Minimac4) ^51^, which generated 65 million genotypes with an imputation score >0.3. In the analyses, we used 45 million genotypes with an imputation score >0.8.

## 4. Data analysis

### 4.1. Bioinformatic QC processing of metagenome sequences

Quality control (QC) of raw sequencing reads was carried out using BBDuk and BBSplit. In the initial QC stage, BBDuk was applied to remove Nextera XT adapter sequences, eliminate PhiX contamination, and perform both quality trimming and filtering. The tool was executed with several key options: ftm=5 to discard any residual bases beyond the 150 bp read length, tbo to trim adapters using paired-end overlap detection, tpe to ensure paired reads were trimmed to equal lengths, qtrim=rl to trim low-quality bases from both the 5′ and 3′ ends, trimq=25 to remove bases below a quality score of 25, and minlen=50 to filter out reads shorter than 50 bp after trimming and contaminant removal. Across samples, 22%–72% of the initial reads were removed at this step, with an average loss of 28%. In the next QC step, reads aligning to the human reference genome (GRCh38.p13) were excluded using BBSplit with default settings, which typically removed fewer than 3% of the BBDuk-processed reads. Subsequently, low-complexity sequences were filtered out using BBDuk’s entropy-based trimming (entropy=0.01), eliminating <2% of the human-depleted reads. This step targeted sequencing artifacts characterized by overrepresented long “G” homopolymers. Overall, our QC workflow mirrored the protocol recommended by HMP24 ^52^, with one modification: because our sequencing approach did not involve PCR amplification, we did not remove duplicate reads, as these could represent true biological signals.

### 4.2. Taxonomic assignment, profiling microbial gene families (KO) and pathways (PTW)

Sequencing reads that passed QC were taxonomically profiled with MetaPhlAn3 ^18^ using its reference database of approximately 1.1 million clade-specific marker genes. MetaPhlAn3 was executed twice for downstream compatibility with different statistical frameworks: (1) relative abundance profiles were produced with default MetaPhlAn3 settings, and (2) a second run was performed using the --unknown-estimation option so that the resulting relative abundances could be converted into count estimates by multiplying by each sample’s total read depth. QC’ed sequences were also profiled for predicted functional elements, i.e., UniRef90 gene-families ^53^ and MetaCyc metabolic pathways ^54^ using HUMAnN3 ^18^ with default parameters. Using the HUMAnN supplied utility tool ‘humann_regroup_table’, the resulting UniRef90 gene-families were regrouped into KEGG Ontology (KO) groups. Pathways and KO groups were filtered to contain only community-level data.

### 4.3 MWAS (Extended Data2-MWAS)

Microbiome-wide association studies (MWAS) were performed consistently across all bioBakery feature types—species, KEGG orthologs (KOs), and pathways—using MaAsLin2 ^55^. Relative abundance values (ranging from 0 to 1) for features present in at least 5% of samples were log-transformed using the “LOG” transformation setting. The three datasets were analyzed using the following three model formulas to account for dataset-specific covariates:

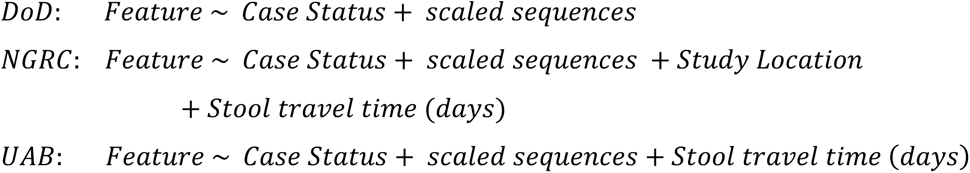

Where “feature” are microbial features (species, KOs, pathways), “case status” corresponds to PD vs NHC, “scaled sequences” is the total sequence count per sample (standardized using ‘scale‘ in R), “study location” corresponds to geographic region where samples were collected, and “stool travel time” is the number of days a stool sample was in transit in US Postal Service. Multiple hypothesis testing was corrected using the Benjamini–Hochberg false discovery rate (FDR) procedure. Conditional analyses were then carried out by including constipation or sex as additional covariates in the models above. Coefficients were extracted for both “case status” and the conditioning covariate.

### 4.4 Meta-analysis (Extended Data2-MWAS)

To generalize per-dataset MWAS results, random effects meta-analyses were applied to bioBakery features that were present in at least two datasets. The MaAsLin2 effect sizes and standard errors were modeled using the *metagen* function in the “meta” R package, specifying method.tau = "REML". Resulting meta-analysis P–values were adjusted with the FDR method. Statistical significance was defined as FDR<0.05 and Cochran’s heterogeneity test P_Het_>0.1.

### 4.5 Networks (Extended Data 5-Networks)

Correlation networks were constructed separately for the PD and NHC microbiomes. Species count tables were used as input for fastspar ^56^ (the C++ implementation of SparCC ^25^) to compute pairwise SparCC correlations, using 100 iterations. To assess significance, permuted P-values were generated by (1) producing 1,000 randomized datasets, (2) recalculating SparCC correlations for each permuted dataset, and (3) determining, for each original correlation, the fraction of permuted correlations with a greater absolute magnitude. Communities of co-occurring species were then identified using the Louvain modularity algorithm ^26^ implemented in the igraph R package (cluster_louvain), applied to correlations meeting the thresholds |r| > 0.2 and uncorrected permuted P < 0.05. For each species, the number of significant correlations (|r| > 0.2 and uncorrected P < 0.05) was quantified using the igraph degree function. For visualization, all significant correlations (|r| > 0.2 and uncorrected permuted P < 0.05) and their corresponding cluster assignments were imported into Gephi. Networks were rendered using the Force Atlas 2 layout algorithm setting non-overlap of nodes with gravity=15 and scaling=5 ^57^, displaying species as nodes colored by significant meta-analysis associations (blue if increased, red if decreased), and edges representing pairwise correlations.

### 4.6 *SNCA* (Extended Data7-*SNCA*)

#### 4.6.1 Selecting PD-associated SNPs at *SNCA*

We used the GWAS Catalog, a database of published GWAS, and searched by selecting “Parkinson disease” and specifying “*SNCA”* in the gene box. Excluding SNPs that did not reach P < 5E-8, the list was reduced to 20 unique PD-associated SNPs annotated to the *SNCA* locus. While cognizant that not all 20 are independent because of linkage disequilibrium (LD), we tested all 20 SNPs and incorporated LD in determining the number of independent tests for the multiple testing Bonferroni correction. LD and correlation between SNPs were calculated twice: once using 1000GENOMES:phase_3:CEU data and the linkage disequilibrium calculator in Ensemble and once using the data from this study to generate study-specific PED/MAP linkage files from PLINK. Results were similar; the latter was plotted in Haploview. SNPs that were in high LD, with D’>0.78 and R2>0.49, were assigned as a haploblock. The 20 SNPs are distributed across a maximum of 9 independent haploblocks. Since MSEA was run on all 20, *N*=9 was used to further Bonferroni-correct the MSEA-generated FDR values, multiplying it by the number of independent tests (FDR*9).

#### 4.6.2 Microbe set enrichment analysis (MSEA)

Testing individual microbes with SNPs would require far greater sample size and power than if microbes could be tested as sets of related species. Thus, we adapted the gene set enrichment analysis (GSEA) framework, substituting microbes (species) for genes, creating microbe set enrichment analysis (MSEA). We used the clusters of fiber degraders and opportunistic pathogens as the microbial sets to test for interaction with *SNCA* SNPs. Fiber degraders and opportunistic pathogens met MSEA requirements as they each had the required minimum of 15 species in the set; the species within a set shared some biological functional relevance, were associated with PD in the same direction (meta-analysis FDR < 0.05, P_Het_ > 0.1), their abundances were correlated with each other, and there was no correlation between fiber degraders and opportunistic pathogens (clusters 9 and 25). The three datasets were pooled to maximize sample size and power, knowing that there is no heterogeneity across datasets in the association of PD with fiber degraders or opportunistic pathogens. To evaluate interactions between the gut microbiome and genetic variants in the *SNCA* locus, we performed association and interaction analyses of individual species and *SNP*s as required for input to conduct MSEA. For each SNP, three models were tested: (1) the association between taxa abundance and SNP dosage in PD cases; (2) the association between taxa abundance and SNP dosage in NHC; (3) the interaction between SNP dosage and PD status (case_status 0|1) on taxa abundance. Because MaAsLin2 does not natively support interaction terms, we manually computed the SNP×PD interaction variable and included it as a fixed effect. For each of the 20 *SNCA* variants selected for analysis, we conducted an MWAS following the same protocol described in **Section 4.3**, with the addition of each variant’s dosage and the interaction term (when applicable). For each tested species, the t-statistic was obtained by dividing the estimated effect size by its standard error. The resulting t-statistics were used as input to the fgsea software for enrichment analysis of two predefined microbe sets: fiber degraders (20 species) and opportunistic pathogens (16 species). MSEA was run using default fgsea parameters. FDR-adjusted P-values were further multiplied by 9 as Bonferroni correction and capped at 1 to correct for the 9 independent haploblocks among the tested variants.

#### 4.6.3 *SNCA*-PD association

Logistic regression was used to quantify the association between 20 SNP alleles of *SNCA* and PD status. Disease state (PD=1, NHC=0) was modeled as the dependent variable. *SNCA* SNP genotype was the primary independent variable and was represented as the continuous imputed allele dosage. The imputed dosage was calculated as the expected number of minor alleles based on the genotype posterior probabilities (dosage = P(G=1) + 2P(G=2)) where G denotes the number of minor alleles (0, 1, 2). Imputed dosage values range continuously from 0 to 2 with 0 corresponding to the homozygous major allele genotype. Minor allele was defined as the less frequent allele in the imputation dataset and used as the effect allele regardless of its direction of association with PD. A linearly additive model was assumed:

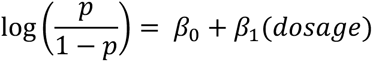

where the coefficient 𝛽_1_ represents the change in log-odds of having PD per one-unit increase in imputed allele dosage, i.e. per additional copy of the minor allele. Odds ratios and 95% confidence interval were obtained by exponentiating 𝛽_1_. No additional covariate adjustment was applied to these models.

#### 4.6.4 Penetrance (Table 4. *SNCA*-Microbe)

We questioned if the risk of PD associated with *SNCA* alleles (i.e., penetrance, measured by odds ratio as a surrogate) is further elevated in people with a dysbiotic feature, namely, high levels of opportunistic pathogens, or low levels of fiber degraders. First, we calculated the mean relative abundance of each dysbiotic feature in NHC and established the upper and lower boundaries of the 95% confidence intervals in NHC. Then we stratified both cases and controls according to the load of the dysbiotic feature in each individual, as follows. Opportunistic pathogens: microbe absent (not detected in the person), present (detected irrespective of relative abundance), and high (relative abundance in the individual was higher than the upper 95% confidence interval boundary in NHC). Fiber degraders: low (relative abundance in subject was lower than the lower 95% CI boundary in controls) and above average (higher than the mean in controls). We then tested the association of the *SNCA* SNP with PD in each stratum, as detailed in **4.6.3 *SNCA*-PD association.**

### 4.7 Subtyping (Extended Data 8-Subtyping)

To classify a metagenome as dysbiotic, we started with the individual-level relative abundances in controls as the reference framework and calculated the mean and 95% confidence interval (CI) of each dysbiotic feature in controls. We then turned to individual-level relative abundances in patients. For features that were elevated in PD, we called a PD gut dysbiotic if the relative abundance of the feature in that person was greater than the upper bound of 95% CI of controls. For features that were reduced in PD, a PD individual’s microbiome was called dysbiotic if the relative abundance of the feature in that person was less than the lower bound of 95% CI of controls. The percentage of patients with a dysbiotic feature was the prevalence of that feature. We recognize that setting the threshold at beyond 95% may be too conservative; it can be adjusted to suit the study. Our working definition of a PD-associated polymicrobial feature was grouping species that (a) were associated with PD in the same direction, elevated or reduced, and, (b) their relative abundances in PD were correlated, they grow together or shrink together, and, (c) they were taxonomically related (*Bifidobacteria* spp, or *Lactobacillus* spp) or had similar biological effects (fiber degraders, or opportunistic pathogens). First, we used the DOD dataset to define six polymicrobial features, estimated their prevalence in DOD patients, and assessed the replication of DOD-defined features in UAB and NGRC. Having established consistency across datasets, we recreated the framework with the pooled data, using MWAS meta-analysis to define dysbiotic features, all 544 NHC to calculate frequency distribution and 95% CI of dysbiotic features, and all 1,006 PD subjects to calculate prevalence of dysbiotic features in PD. We ran the analysis once for dysbiotic clusters and once for selected species.

To estimate the effect of prescreening before enrolment to the clinical cohort, we used parameters used in a completed trial (https://clinicaltrials.gov/study/NCT04658186), namely 30% improvement in treatment, measured as a placebo effect in the untreated arm being 2.285 increase in UPDRS I-III score/year, and 30% improvement in the treatment arm being (2.285*1-0.3)=1.59 increase in UPDRS I-III score/year. We calculated and plotted power and sample size, setting ⍺=0.1, treatment effect=30%, for four scenarios: prescreened and selected for having the dysbiotic feature, not prescreened with 70%, 30%, and 10% of the cohort having the dysbiotic feature.

## 5. Functional inference

### 5.1 Functional inference on dysbiotic features of PD (**Figure 2**. Genes and pathways)

KEGG ortholog gene families (KO) and pathways were profiled as in **Section 4.2** and analyzed as in **Sections 4.3 and 4.4**. KO and pathways whose relative abundances were significantly elevated or reduced in PD vs. NHC in meta-analysis (FDR<0.05, P_Het_>0.1) were considered for functional inference. Species that contributed to the KO and pathways were used to trace the origin of the altered feature. Contributing species were extracted for each of the three datasets, as species do not enter the meta-analysis of KO and pathways, and there is no aggregate compilation of contributing species. Contributing species were available for PD and NHC separately per dataset. We used the PD cohort as the source of contributing species if the KO or pathway was elevated in PD and used NHC as the source if the KO or pathway was elevated in NHC.

### 5.2 Functional inference for microbiome-associated *SNCA* SNPs (Figure 3. *SNCA*)

SNPs in 4 haploblocks showed significant association with opportunistic pathogens or fiber degraders. SNPs within each haploblock yielded nearly identical results for association with microbiome, as expected due to their strong LD. Thus, any one SNP from each haploblock could be the representative for that block. Databases like GTEx, which assemble genome-wide data on regulatory SNPs, often include one SNP per LD block. Screening GTEx, we found one SNP for each of the 4 haploblocks, and they were associated with differential expression and splicing of *SNCA*. We used those four SNPs in the UCSD Genome Browser to map their position on the chromosome and overlay with regulatory sequences, including enhancers, transcription factor binding sites, and the *SNCA* promoter.

## Supporting information

Extended Data 1-STORMS

Extended Data 2-MWAS

Extended Data 3-Constipation

Extended Data 4-Sex

Extended Data 5-Networks

Extended Data 6-Inference

Extended Data 7-SNCA

Extended Data 8-Subtyping

## Data and code availability

All data used and generated in this study are publicly available. The raw, unprocessed metagenomic sequences and their associated subject characteristic metadata were deposited in the NCBI Sequence Read Archive (SRA) under BioProject ID PRJNA1454017 (https://dataview.ncbi.nlm.nih.gov/object/PRJNA1454017), and SRA has completed Human Contamination Screening to remove human DNA that may be a subject identifier. Post-QC and taxonomic and functionally profiled sequences and the subject characteristic metadata are in the “Source Data” document on Zenodo [https://zenodo.org/records/19265067]. Genome-wide genotypes are on the NCBI Database of Genotypes and Phenotypes (dbGAP) study accession phs000196 [https://www.ncbi.nlm.nih.gov/projects/gap/cgi-bin/study.cgi?study_id=phs000196]. The code used to perform sequence QC, taxonomic and functional profiling, statistical analyses, and figure generation is on Zenodo [https://zenodo.org/records/19265067]. The data, code, protocols, and key lab materials used and generated in this study are listed in a Key Resources Table alongside their persistent identifiers in **Table 5. Key Resources**. For the purpose of open access, the authors have applied a CC BY public copyright license to all Author Accepted Manuscripts arising from this submission.

## Reproducibility

The results can be reproduced using either the raw sequence (on SRA) or processed sequences (on Zenodo), together with the metadata (on SRA and in “Source Data” on Zenodo), the code (on Zenodo), and following the Methods section of the manuscript.

## Author contribution

All authors have made a substantial contribution to the work. They have all read and agree with the manuscript’s content and authorship. All authors have agreed to be personally accountable for their own contributions. Every author agrees that if any questions arise regarding the accuracy or integrity of any part of the work, even those in which the author was not personally involved, they will ensure that the questions are appropriately investigated, resolved, and the resolution documented in the literature.

## Acknowledgements

We sincerely thank the individuals with Parkinson’s disease, their spouses and caregivers, and the healthy volunteers who participated in this research project.

